# Structural reorganization underlying stress-induced cytoplasmic solidification in yeast

**DOI:** 10.64898/2025.12.22.696049

**Authors:** Sara Katrin Goetz, Marie-Christin Spindler, Rasmus Kjeldsen Jensen, Willram Scholz, Evgenia Zagoriy, Julia Mahamid

## Abstract

Cells employ diverse strategies to rapidly adapt to sudden environmental changes. In yeast, cytoprotective solidification in response to starvation and energy depletion (ED) has been reported and associated with extensive mesoscale macromolecular assembly. Yet, the structural and molecular basis underlying such whole-cell level liquid-to-solid phase transitions remain unknown. Here, we use cryo-electron tomography to characterize the subcellular organization of intact yeast cells exposed to ED and other stressors, and to untangle the effects of molecular crowding versus cytoplasmic acidification previously suggested to underpin solidification. We visualize self-assembly of macromolecules and complexes into ordered assemblies and condensates under ED, and quantify ribosome and polysomes concentrations to probe changes in cytoplasmic crowding. Combined with live-cell microscopy, we pinpoint supramolecular assembly induced by acidification, rather than a uniform increase in intracellular crowding, as the structural basis of cytoplasmic solidification that supports yeast cells’ adaptation in response to environmental stresses.

## INTRODUCTION

Organisms are constantly challenged by a range of stressors. In order to survive, cells employ a set of common stress response mechanisms^1–8^. Stress responses are extensively studied in yeast^9^, which adapt their metabolism, growth rate and morphology when challenged with stressors that include heat^10,11^, osmotic and oxidative stresses^12,13^. When faced with nutrient scarcity, yeast can further transition into a reversible dormant state that is characterized by low metabolic activity and is reported to affect the whole proteome in both *S. cerevisiae* and the distantly related *S. pombe*^14,15^. Indeed, glucose depletion leads to a global reduction in transcription, mRNA degradation^16^, and translation^17,18^, accompanied by alteration in autophagy^19–21^. Within minutes of glucose or acute energy depletion (ED; simultaneous inhibition of glycolysis and mitochondrial respiration), intracellular ATP is almost completely consumed^22–24^.

Strikingly, the cytosol of both *S. cerevisiae* and *S. pombe* is reported to solidify in response to glucose deprivation and ED^25–27^. Thus, beyond responses at the molecular scale, survival under unfavourable conditions is associated with alterations to the physical properties of the cell, attributed in part to changes on the mesoscale^28^. In recent years, the formation of supramolecular assemblies and biomolecular condensation have been proposed as mechanisms that allow cells to promptly reorganize their existing proteomes to promote survival^7,29–34^. In contrast to irreversible aggregation, regulated assembly of macromolecules at the mesoscale provides a protective mechanism^27^. On the molecular level, assembly can regulate the conformational landscape of the underlying proteins, which in turn, can modulate their catalytic activities^8,35–39^. Indeed, early genome-wide light microscopy screens in *S. cerevisiae* identified more than 200 proteins to form cytosolic foci or rods upon glucose starvation^29,35,40–43^, including essential metabolic enzymes, such as fatty acid synthase^44^. More recent cryogenic electron microscopy and tomography (cryo-EM/ET) studies directly link conformational changes upon polymerization of the enzymes CTP synthetase, Glutamine synthetase, Acetyl-CoA synthetase (Asc1) and TORC1, to regulatory mechanisms of their activity^36,38,39,45^.

However, what triggers mesoscale assembly upon nutrient deprivation in yeast and what structural changes amount to their documented global cytoplasmic solidification remain unclear. Starvation and ED are accompanied both by an increase in molecular crowding, inferred from an approximate 7% reduction in cell volume and in the mobility of organelles, endogenous macromolecules and foreign tracer particles, and by intracellular acidification^25,26,46,47^. Both crowding and acidification are common responses in yeast exposed to many stresses, including osmotic stress and heat shock, and are therefore hypothesized to be potential drivers of cytoplasmic solidification^25,26,48,49^.

Changes in intracellular crowding affect many biophysical properties, including intracellular viscosity and protein diffusion, folding, conformations, interactions, oligomerization, and condensation^50–57^. For example, a controlled increase in intracellular crowding induces condensation of the synthetic SUMO_10_ and SIM_6_ system^58^ in yeast and human cells^59^. Consequently, changes in molecular crowding are suggested to be key in modulating the biophysical properties of the yeast cytoplasm. In the cytoplasm, ribosomes and polysomes represent major crowders owing to their large size and high abundance. Polysomes disassemble within 10 minutes of ED and 30 minutes of glucose starvation ^17,47,60^ following downregulation of protein translation^37^. Polysome disassembly immediately after nutrient depletion was indeed recently shown to transiently fluidize the cytoplasm^47^. But polysomes partially reform after 60 minutes of starvation, likely for (longer term) adaptation^61^. Over these longer periods of glucose deprivation and under ED, particle mobilities are also profoundly reduced^25,26^. Thus, polysomes may play a role as polymeric crowders in inducing the liquid-to-solid transition of the yeast cytoplasm at late stages of nutrient stress^25^.

At the same time, the significant reduction in ATP levels under starvation and ED^17,21,23,24^ leads to disassembly of the vacuolar-ATPase^62^. Under nutrient-rich conditions, the pump continuously removes protons from the cell into the vacuole to maintain neutral cytosolic pH^63,64^. V-ATPase disassembly leads to a breakdown of the proton gradient^65,66^ and to cytosolic acidification^26^. Since the isoelectric points of a large fraction of the yeast proteome falls within the acidic range, their net charge is reduced at a low intracellular pH^67^. Reduced protein net charge, in turn, is shown to promote intramolecular interactions and the formation of biomolecular condensates^26,68–71^. Thus, acidification is suggested to promote large-scale supramolecular assembly that could contribute to apparent cytoplasmic solidification.

Understanding the structural and molecular underpinnings of subcellular reorganization under prevalent stress conditions can shed light on fundamental physicochemical mechanisms that promote cell survival. Here, we combined cryo-ET of intact yeast cells with live-cell light microscopy to map changes in macromolecular structure, distribution and assembly under ED, osmotic stress and in response to ectopic acidification. We determined how these stressors affect intracellular crowding and cytosolic pH in the two evolutionary distant yeast species, *S. pombe* and *S. cerevisiae*. Our data quantitatively assess the cellular and structural changes under different stress conditions and pinpoint cytosolic acidification as a general stress signal that drives large-scale supramolecular assembly.

## RESULTS

### Cryo-ET reveals large-scale intracellular reorganization in energy depletion

To visualize and characterize changes that accompany yeast cell solidification upon ED^25,26^ at the subcellular and macromolecular level, we acquired more than 600 label-free cryo-electron tomograms of wild-type *S. pombe* and *S. cerevisiae* cells (Table 1, Figure 1A, B, Figure S1, Methods).

**Table 1.** Summary of the cryo-ET data analysed for the two yeast strains under native state and different stress conditions. A quantitative assessment of the appearance of the different supramolecular assemblies in the data is provided. ED: energy depletion, GD: glucose depletion, OS: osmotic stress, Mito: mitochondrion.

| Sample | #tomos | #tomos<br>with<br>Cytosolic<br>Filaments | #tomos<br>with<br>Nuclear<br>Filaments | %tomos<br>with<br>Filaments | #tomos<br>with<br>Mito<br>lattice | %tomos<br>with<br>Mito<br>lattice | #tomos<br>with<br>FAS<br>assembly | %tomos<br>with<br>FAS<br>assembly |
| --- | --- | --- | --- | --- | --- | --- | --- | --- |
| *S. pombe* |  |  |  |  |  |  |  |  |
| native state | 48 | 0 | 0 | 0 | 0 | 0 | 0 | 0 |
| 1 h ED | 117 | 36 | 3 | 33 | 26 | 22 | 43 | 37 |
| 3.5 h ED | 76 | 25 | 1 | 34 | 18 | 24 | 53 | 70 |
| 17 h ED | 34 | 11 | 1 | 35 | 6 | 18 | 9 | 26 |
| 4 d GD | 31 | 13 | 1 | 45 | 0 | 0 | 0 | 0 |
| 15 min OS | 43 | 3 | 1 | 7 | 1 | 2 | 0 | 0 |
| 8 min pH 4.5 | 10 | 2 | 0 | 20 | 3 | 30 | 8 | 80 |
| 50 min pH 4.5 | 66 | 19 | 0 | 28 | 12 | 18 | 40 | 61 |
| *S. cerevisiae* |  |  |  |  |  |  |  |  |
| native state | 64 | 0 | 0 | 0 | 0 | 0 | 0 | 0 |
| 1 h ED | 62 | 11 | 3 | 23 | 0 | 0 | 10 | 16 |
| 6 h ED | 56 | 7 | 3 | 18 | 0 | 0 | 8 | 14 |
| 10 min OS | 22 | 0 | 0 | 0 | 0 | 0 | 0 | 0 |
| 35 min pH 4.5 | 13 | - | - | - | - | - | - | - |

**Figure 1.**
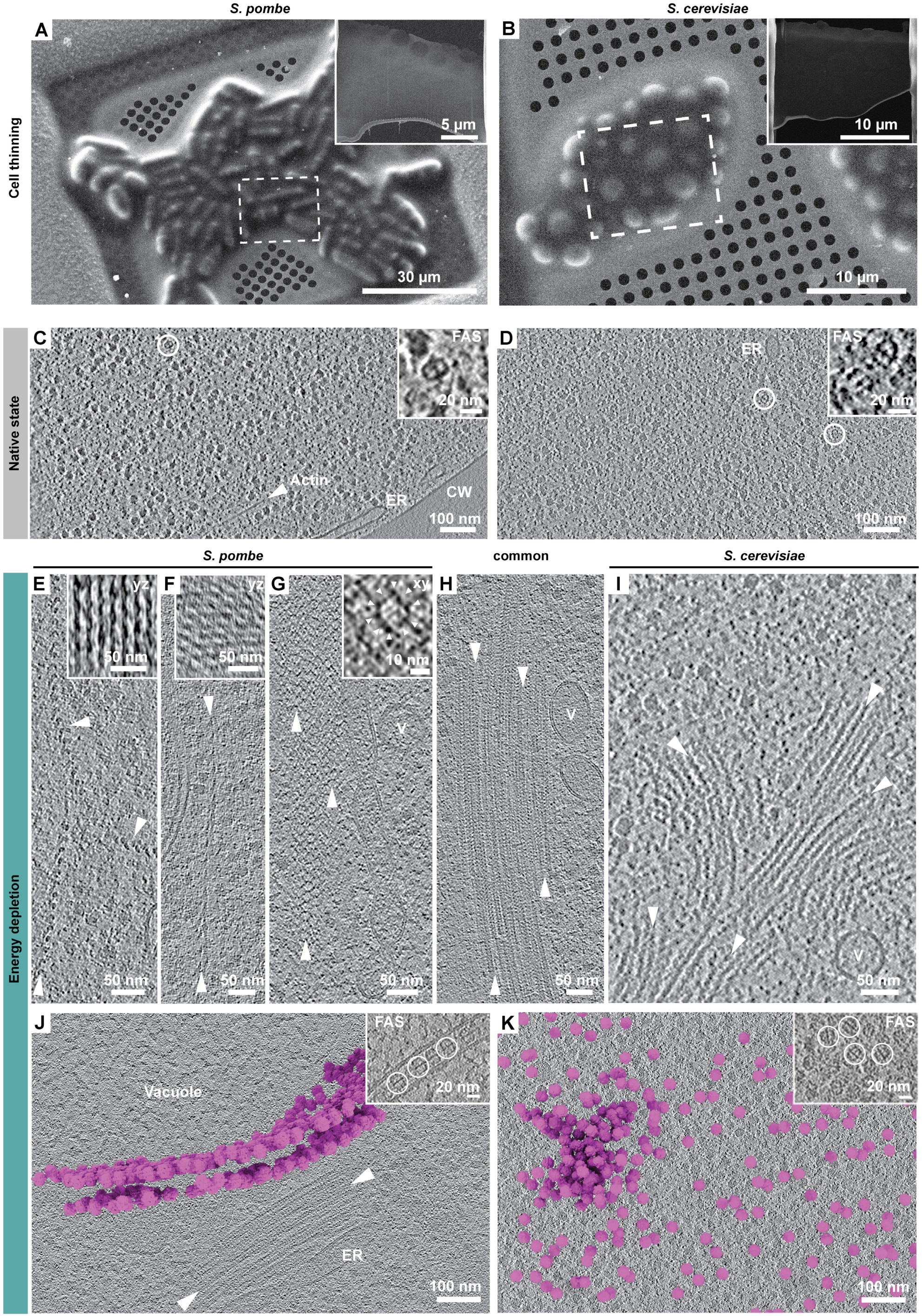
Cryo-ET of yeast cells under normal nutrient conditions and energy depletion reveals structural reorganization of the cytoplasm. (A, B) Scanning electron microscopy (SEM) images of plunge-frozen *S. pombe* (A) and *S. cerevisiae* (B) cells. Insets: SEM view of thin cellular slices (lamellae) generated by cryo-focused-ion beam (FIB) milling of the areas targeted in A and B (dashed box). (C, D) Tomographic slices of native state *S. pombe* (C) and *S. cerevisiae* (D), depicting the cytosol crowded with ribosomes (abundant dark particles), barrel shaped fatty acid synthase (FAS) complex (white circles, and enlarged in insets), an actin filament (arrowhead), the endoplasmic reticulum (ER), and the cell wall (CW). (E-K) Tomographic slices at 1h ED show macromolecular assemblies with different (polymeric) architectures (white arrowheads): (E-G) In *S. pombe*, two filamentous assemblies with a pitch of ∼30 nm and 33 nm (E, F insets: enlarged orthogonal views), and a lattice composed of putative 10 subunits (G: enlarged in inset). V: vesicle. (H) A common lattice architecture found in both yeast species. (I) A filamentous cytosolic assembly specific to *S. cerevisiae*. (J, K) FAS (annotated in magenta) partitions into mesoscale assemblies in a species-specific manner under ED. Insets: enlarged views; individual FAS complexes encircled. Arrowheads in J indicate adjacent filamentous assembly.

Tomograms from cells under normal nutrient conditions (native state) showed the cytosol containing ribosomes, barrel-shaped fatty acid synthase (FAS) complexes, cytoskeletal actin filaments, and membrane-bound organelles (Figure 1C, D). At 1h ED, tomograms revealed changes to the morphologies of organelles, most prominently to mitochondria, lipid droplets and vacuoles (Figure S2): mitochondria are tubular under native conditions^72^, but appeared spherical in ED cells (Figure S2A, B).

We observed events of mitochondrial fission (Figure S2B), which had been reported for *S. pombe* under glucose starvation^73,74^. The outer mitochondrial membrane of ED *S. pombe* was decorated with ribosomes (Figure S2A) in line with recent reports of mitochondria-associated hibernating ribosomes in starved *S. pombe* cells^75^, and further exhibited surface lattices (Figure S2A) whose structural signature resembled assemblies of the GTPase dynamin related protein Dnm1^76–78^ required for mitochondrion fission. Lipid droplets (LDs) in ED were frequently engulfed by or connected to vacuoles, and occasionally to the nuclear envelope (Figure S2C-H), reproducing reports of LD fragmentation and membrane invagination in ED or starved *S. cerevisiae*^27,79^. Our cryo-ET data further revealed crystalline layers in LDs (Figure S2C, D) that result from phase transitions of sterol lipid^80,81^. Our observations of morphological changes to cellular organelles in ED thus overall align with previous studies.

Beyond changes to organelles, we observed a number of structurally-distinct molecular assemblies under ED that we visually assigned to five different classes of filament or lattice architectures (Figure 1E-I). Three of the oligomeric structures were unique to *S. pombe* (Figure 1E-G), one was common to both yeast species (Figure 1H) and one specific to *S. cerevisiae* (Figure 1I). Their dimensions and structural signatures differed from those of known cytoskeletal components (i.e., actin filaments and microtubules). One of the polymers observed in *S. pombe* (Figure 1E) and the oligomer unique to *S. cerevisiae* (Figure 1I) were observed in both cytosol and nucleus. Previous light microscopy studies^38,40–42^ identify a number of proteins that form foci and rods upon nutrient deprivation, which qualify as candidates for the filamentous assemblies observed in our study. However, structural data is available only for a small subset of these candidates in their assembled, filamentous state. Recently, Hugener et al. identified filaments of Acs1 in the nucleus and cytoplasm of meiotic yeast with a diameter of 13 nm^45^. In 2020, Marini et al. demonstrated that eIF2B forms filament bundles in ED *S. cerevisiae* using room-temperature correlative light and electron microscopy (CLEM)^27^. Subtomogram averages of these helical filaments revealed a diameter of 16 nm and a repeat of 14.5 nm. These dimensions roughly match the 13 nm repeat and 13 nm diameter that we measured for the oligomer unique to *S. cerevisiae* (Figure 1I). Based on the similarity of the dimensions and morphology, we speculate that eIF2B could form the basis of this oligomeric structure. However, to unambiguously identify the proteins that form the diversity of the structures we observed upon ED, and to decipher the functional outcomes of assembly, high-resolution structures, in combination with cryogenic CLEM approaches, will be required^82^.

With a volume occupancy of ∼5%, the structured assemblies collectively make up a considerable fraction of the cytosolic volume (Methods). In addition, ribosome excluded areas indicated the presence of less ordered molecular assemblies, potentially condensates, which we could not further characterize due to the lack of a defined structural signature. Tomograms acquired at later time points of ED indicated that the assemblies grow over time (*S. pombe*: 3.5h and 17h, *S. cerevisiae*: 6h; Figure S3A-E), suggesting that while the cells macroscopically solidify within one hour of ED^25,26^, cytosolic rearrangement continues for several hours. We further examined whether structured assemblies form in response to more physiological starvation conditions, and indeed observed large-scale lattice-like structures after four days of glucose deprivation (Figure S3F).

Finally, we visualized mesoscale assembly of FAS which interestingly exhibited species-specific architectures (Figure 1J, K). The energy-dependent fungal type I FAS is an essential metabolic multi-enzyme complex that catalyzes the synthesis of long-chain fatty acids. These are required to build membranes, serve as precursors of second messenger molecules, and to store energy in the form of triacyl glycerides (TAGs) in LDs^83^. Under ED, FAS arranged into 2D monolayers associated with adjacent membranes in *S. pombe* (Figure 1J), but formed seemingly randomly organized clusters in *S. cerevisiae* (Figure 1K). A previous light microscopy study showed that FAS evenly distributes within the cytosol under native conditions, but condenses into foci after several days of glucose starvation in *S. cerevisiae*^44^. Room temperature CLEM localized these FAS foci to ribosome-excluded areas, and purified complexes from this condition were found to be fully assembled^44^. The enzyme’s functional state and its structural arrangement within the cytoplasmic foci, however, remained unknown. FAS being a large macromolecular complex (2.6 mDa) with a characteristic shape makes it readily identifiable in cryo-ET. It thus presented a suitable system to interrogate the properties of mesoscale assemblies formed by structured macromolecular complexes in response to stress.

### Species-specific differences in the architecture and dynamics of FAS assemblies

To probe whether the transition of FAS from individual, dispersed complexes (Figure 1C, B) into mesoscale assemblies (Figure 1J, K) induces structural changes that affect its enzymatic activity, we generated subtomogram averages of the complex localized in *S. pombe* and *S. cerevisiae* under native state conditions and 1h ED. We then performed focused classifications on the catalytic sites of the resulting maps to analyze potential functional differences (Figure 2A, Methods). The low particle numbers of the dispersed FAS complexes in native state cells lead to maps resolved at 27 Å and 32 for *S. pombe* and *S. cerevisiae,* respectively (Figure 2A, Figure S4A). Under ED, we resolved FAS to 13 Å for *S. pombe* and 17 Å for *S. cerevisiae* (Figure 2A, Figure S4A).

**Figure 2.**
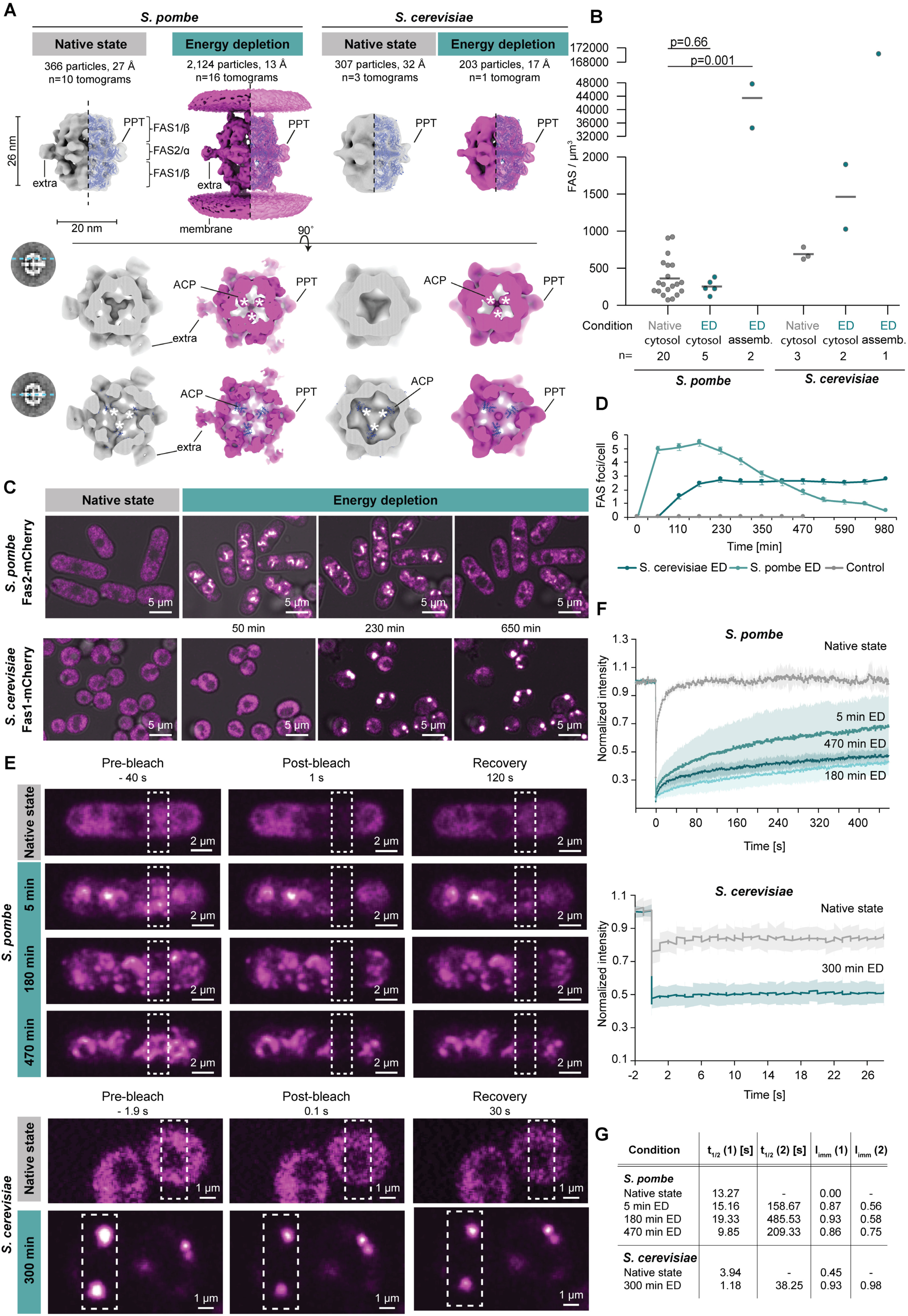
Nutrient and species-dependent differences in FAS conformation and assembly. (A) FAS subtomogram averages refined with D_3_ symmetry from native state (grey) and 1h ED (magenta) cells. Tomogram and particle numbers, and global resolutions are indicated. The in-cell averages are fitted with the published *S. cerevisiae* structure (PDB 2uv8^84^, blue). Cross-sectional views (bottom rows) show acyl carrier proteins (ACPs, white asterisks) close to the α-wheel in the native state, but at the top of each half dome in ED cells. (B) FAS local concentrations calculated based on particle numbers per cellular volume determined from complexes localized in native state and ED tomograms. Statistical significance (p<0.05) evaluated with Kolmogorov–Smirnov test. Number of analysed tomograms (n) is indicated for each condition. Significance was not evaluated statistically for *S. cerevisiae* due to the small dataset included in the analysis. (C) Dynamics of FAS assembly upon ED. In *S. pombe*, FAS-mCherry forms foci immediately upon ED, while assembly is delayed in ED *S. cerevisiae*. (D) Corresponding quantification of FAS foci counts per cell. Data are shown as mean ± SD. Number of cells: *S. cerevisiae* control n = 52 and ED n = 92; *S. pombe* control n = 22 and ED n = 58. (E) FRAP (dashed boxes) of FAS-mCherry-expressing *S. pombe* (top) and *S. cerevisiae* (bottom) in native nutrient state and at different timepoints of ED. (F) Corresponding FRAP curves. Normalized fluorescence recovery curves (solid lines) ± SD (shaded area) from 5 and 21 replicates per condition for *S. pombe* and *S. cerevisiae*, respectively for each time point. (G) Half times (t_1/2_) and immobile fractions (I_imm_) quantified from FRAP for free FAS and FAS foci. Recovery curves were fitted with a one-step (native state) or two-step exponential model (ED) accounting for cytosolic only (native state) or cytosolic and condensed protein complexes (ED). The overall topology of all four maps matched the published crystal structure of purified *S. cerevisiae* FAS (PDB: 2uv8^84^), showing three FAS1/ß subunits forming each of the two half domes connected through a central α-wheel comprised of six FAS2/α subunits. At the plane of the α-wheel, we resolved two different densities. One density fitted the C-terminal phosphopantetheinyl transferase (PPT) domain (Figure 2A). In the presence of ATP, the PPT domain attaches coenzyme A to the acyl carrier proteins (ACP) and thereby activates the synthesis of fatty acids^85,86^. The other density, observed only in *S. pombe,* connected to the N-terminus of FAS2/α and could not be assigned by fitting published structures. While we found FAS to be fully assembled in both native state and ED cells, the maps revealed marked differences in the catalytic sites. In the native state, three ACPs fit into densities located close to the α-wheel (Figure 2A). In this conformation, the protein is suggested to shuttle the nascent acyl chain between the different catalytic sites of the complex in order to synthesize long chain fatty acids, indicating an active conformation^85^. Under ED, these densities were located at the top of each half dome, similar to the rotated conformation described for low levels of NADPH^86^. This conformation has been linked to a stalled, hibernating state of the complex^86^ (Figure 2A). Thus, the fully assembled FAS complexes under ED are likely catalytically inactive.

The subtomogram averages further reflected the species-specific differences in FAS organization within the higher-order assemblies observed under ED (Figure 1J, K). In *S. pombe*, three unassigned densities on top of each half dome connected the complex to the adjacent membranes. The identity of the connecting densities remains elusive as they could not be identified by fitting of published structures. The sequence identity for FAS1/ß and FAS2/α between *S. pombe* and *S. cerevisiae* is 48% and 59%, respectively. Fitting of the *S. cerevisiae* FAS complex crystal structure into the *S. pombe* map revealed that two unstructured loops, containing large, non-conserved residues (K453, V470 and K497) insert into the unassigned density and potentially form the interaction interface with an unknown partner (Figure S4B-D). Furthermore, the *S. pombe* unidentified densities located at the level of the α-wheel (alternating with the PPT) did not link neighboring complexes, suggesting that complexes within the monolayers either do not directly interact or that putative linkers are too flexible to be resolved. Our efforts to generate proteomics data to assist identification of FAS interaction partners in ED failed due to the difficulty in lysing the solidified ED cells. In the *S. cerevisiae* average, the membrane connecting densities were absent, in line with the assembly of FAS complexes in a seemingly random organization (Figure 2A, Figure 1K).

Quantification of the local concentrations of FAS complexes per cytosolic volume from the cellular cryo-ET data (Figure 2B, Methods) revealed a significant increase of 120-fold for *S. pombe* (43,423 ± 7,727 particles/μm^3^) and 250-fold for *S. cerevisiae* (170,332 particles/μm^3^) in the 1h ED assemblies compared to the dispersed complexes in the native state (363 ± 255 particles/μm^3^ for *S. pombe* and 690 ± 84 particles/µm^3^ for *S. cerevisiae*). In ED *S. pombe*, the concentration of free FAS complexes (not engaged in the assemblies) was similar to the native state. In *S. cerevisiae*, the concentration of FAS complexes in the vicinity of assemblies was higher by more than two-fold compared to normal nutrient conditions (Figure 2B).

The above-described species-specific differences in FAS assembly architectures suggested possible differences in assembly dynamics of the complex. We thus acquired live-cell confocal fluorescence microscopy data of endogenously tagged FAS (FAS-mCherry) to monitor the formation of assemblies under ED in the two species over time (Figure 2C, Methods). Under native conditions, FAS signal was evenly dispersed in the cytosol (Figure 2C). At the onset of ED, the FAS signal concentrated into distinct foci. In *S. pombe*, the foci exhibited half-moon shapes in agreement with our observations of FAS assembly along curved membranes, i.e., vacuoles, by cryo-ET. The number of FAS foci reached a maximum at around 3.5 h of ED, and dropped steadily thereafter. Assembly dynamics were delayed in *S. cerevisiae*, as foci only started to form around 1h into ED and reached a plateau with 2-3 foci per cell after around 3h (Figure 2D). In order to discern whether the nutrient-dependent FAS assemblies display liquid-like or solid properties, we probed their mobility using fluorescence recovery after photobleaching (FRAP) (Figure 2E-G). Under native conditions, the dispersed FAS freely diffused in the cytosol with a recovery half time of around 13 and 4 seconds for *S. pombe* and *S. cerevisiae,* respectively. Upon ED, the mobility of the complex decreased drastically both in the cytosolic and condensed fractions. In *S. pombe*, FAS dynamics slowed down within 10 minutes of ED with immobile fractions of roughly 90% and 60% for assembled and free FAS, supporting solidification of the cells. Recovery dynamics decreased even further after 3h and appeared to slightly increase only beyond 8h of ED (Figure 2F, G), in agreement with the observed dissolution of foci in light microscopy movies (Figures 2C) and the less frequent observation of FAS assemblies by cryo-ET at later time points (Figure S3). For *S. cerevisiae*, FRAP of FAS-mCherry was assessed at 6h of ED when foci formation had reached a plateau. Here too, the formation of FAS foci coincided with a significant decrease in the mobility of the complex, reflected in an increase of the immobile fraction from an average of 45% in the control to more than 90% beyond 5h of ED (Figure 2G).

In summary, the decrease in FAS mobility in both species, together with our characterization of the nano-scale changes in particle distributions, identify FAS assembly under ED as an example for nutrient-dependent macromolecular de-mixing. Importantly, we identified that the metabolic complex adopts a likely catalytically inactive state under this stress condition. We next aimed to investigate which of the factors previously suggested to contribute to cytosolic solidification, *i.e.* molecular crowding and acidification, correlate with the appearance of such molecular assembly/de-mixing.

### Marginal changes in molecular crowding occur under ED

A reduction in cell volume of around 7% has been reported for ED yeast^13,14^, which suggests an increase in intracellular crowding. It is further proposed that increased molecular crowding plays a role both in the induction of biomolecular condensation through entropic effects associated with volume exclusion^87^ and in the global liquid-to-solid phase transition of the yeast cytosol^25^.

To probe nutrient-dependent changes in crowding at the subcellular level, we determined ribosome concentrations based on cryo-electron tomograms of native state and 1h ED cells (Figure 3A, B, Methods, Figure S5A). Ribosomes are megadalton-scale complexes, highly abundant and occupy a large volume fraction of the cytosol (around 20%)^50^. Their local concentrations can tune the effective concentration of other biomolecules by modulating the available physical space.

**Figure 3.**
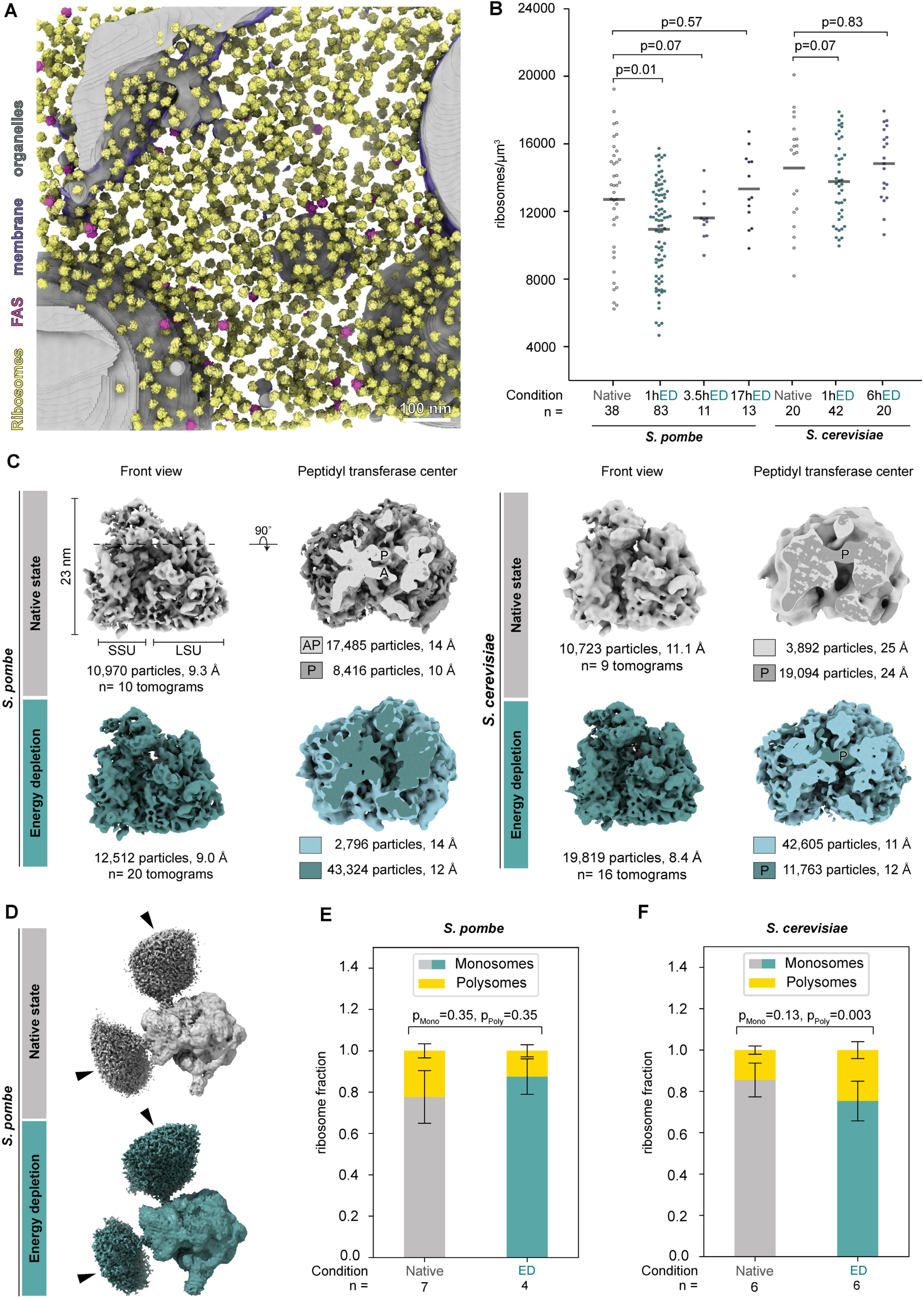
Molecular crowding probed by ribosome concentrations is not significantly altered under different nutrient conditions. (A) Example of comprehensive 3D annotation of a native state *S. pombe* tomogram with ribosomes (yellow), FAS complexes (magenta), membranes (purple) and organelles (grey). (B) Ribosome concentrations calculated based on particle numbers per cellular volume determined from tomograms of native state and different time point of ED. Statistical significance (p<0.05) evaluated with Kolmogorov–Smirnov test. Number of analysed tomograms (n) is indicated for each condition. (C) Nutrient-dependent states of the yeast 80S ribosomes with different occupancies of A- and P-site tRNAs at the peptidyl transferase center. Tomogram and particle numbers, and determined resolutions are indicated. SSU and LSU: small and large ribosomal subunits. (D) Ribosome averages showed adjacent densities (arrowheads) indicative of neighboring ribosomes in a polysome configuration in all nutrient states. (E, F) Quantification of monosome and polysome fractions in native state and 1h ED *S. pombe* (E) and *S. cerevisiae* (F) tomograms. Statistical significance (p<0.05; B, E, F) performed with Kolmogorov–Smirnov test.

Ribosome concentrations in the cytosol of native state cells amounted to an average of 5.7×10^5^ ribosomes/cell for *S. pombe* and 2.4×10^5^ ribosomes/cell for *S. cerevisiae* (Figure 3B, Methods), in line with previous proteomics and cryo-ET data available for *S. pombe*^14,88^ and *S. cerevisiae*^59,89–93,94^. In contrast to an expected increase in molecular crowding, ribosome concentrations in 1h ED cells were similar to native state cells: in *S. pombe*, we determined an average of 12,708 ± 3,460 ribosomes/µm^3^ in native state and 10,840 ± 2,926 ribosomes/µm^3^ in 1h ED. In *S. cerevisiae*, we detected an average of 14,517 ± 3,193 ribosomes/µm^3^ in native state and of 13,768 ± 2,379 ribosomes/µm^3^ in 1h ED. Ribosome concentrations did not change significantly in cells that were exposed to longer ED (*S. pombe*: 3.5h and 17h ED, *S. cerevisiae*: 6h ED; Figure 3B). Thus, our measured ribosome concentrations do not support the hypothesis wherein a global increase in molecular crowding upon ED leads to a reduction of particle diffusion dynamics that characterize yeast cell solidification.

Nutrient deprivation is known to lead to a global downregulation in the translational activity of ribosomes^17,60,61^, and to the disassembly of polysomes^17,46,47,95,96^, which are expected to contribute significantly to intracellular crowding. To assess these aspects, we first generated subtomogram averages from all detected cytosolic 80S ribosomes (Figure 3C, Methods). In native state cells, we obtained consensus maps with resolutions of 9.3 Å for *S. pombe* and 11.1 Å for *S. cerevisiae* (Figure 3C, Figure S5B, C). For 1h ED, we obtained maps with resolutions of 9.0 Å for *S. pombe* and 8.4 Å for *S. cerevisiae* (Figure 3C, Figure S5B, C). All four maps resembled the published *S. cerevisiae* ribosome structure and subtomogram average (PDB: 6tnu^97^ and EMDB: 4372^59^). We detected only fully assembled ribosomes, in contrast to non-translating free large ribosomal subunits. Next, to determine the ribosome functional state, we performed focused classifications on the peptidyl-transferase center (PTC) and analyzed densities at binding sites of translation factors and tRNAs (Figure 3C, Methods). For each condition, we identified a major and a minor class: ribosomes were overall translationally active in the native state in both yeast species indicated by the P-site being occupied with tRNA density in most ribosomes^98^. Specifically, the major class in *S. pombe* (67.5% of particles) exhibited tRNA densities both at the A- and the P-site. In the minor class (32.5%), a tRNA density at the P-site and an amino acyl-tRNA density located at the PTC entry site were observed. In *S. cerevisiae,* the major class also contained a tRNA density at the P-site, while 16.9% of particles showed an empty PTC, likely representing non-translating complexes. The maps from 1h ED revealed mostly empty PTCs (Figure 3C), representing fully assembled, but translationally inactive ribosomes. In *S. pombe*, a density was resolved at the tRNA entry site. Given the moderate resolution of the map, this density could represent an amino-acyl tRNA density, but it may also accommodate eRF1 (PDB 3jah^99^), a release factor that terminates protein synthesis in eukaryotes^99–102^ or eEF2 (PDB 4v4b^103^), a eukaryotic elongation factor that facilitates GTP-dependent ribosome translocation and is inhibited by the EF-2 kinase leading to inhibition of protein synthesis^104^. In *S. cerevisiae*, the PTC was empty in nearly 80% of ribosomes while the remaining particles contained a tRNA density at the P-site (Figure 3C). The latter hints at a subset of ribosomes actively translating stress response-related proteins upon 1h ED, in line with previous reports of polysome reformation after 60 minutes of glucose starvation^61^. Thus, we show that ED overall leads to downregulation of protein synthesis.

Interestingly, despite the fact that our focused classifications on the PTCs suggest ribosomes to be largely translationally inactive at 1h ED, both *S.pombe* and *S. cerevisiae* ribosome maps showed densities indicative of neighboring ribosomes at positions that match their engagement in polysomes (similar to native state; shown for *S. pombe* in Figure 3D). We therefore next quantitatively assessed whether polysomes persist under ED conditions and may contribute to the reported reduction in particle mobility^25,26^ due to their suggested role as polymeric crowders^46,47^. For this purpose, we used the refined positions and orientation of ribosomes from a subset of tomograms that were comprehensively manually curated to achieve an estimated 90-95% coverage of ribosomes (Methods). We defined polysomes and quantified their numbers in native state and ED cells (Figure 3E, F, Figure SA, B, Methods). Ribosomes were considered to be part of a polysome when the distance between the mRNA exit and entry site of two neighboring ribosomes was below 8 nm^46,105^ (Figure S6). Otherwise, they were assigned to the monosome fraction. In *S. pombe*, a decrease in polysome fraction from 22.3% in native state cells to 12.4% in 1h ED was non-significant (Figure 3E; p>0.5). In contrast, the polysome fraction increased in *S. cerevisiae* upon ED from 14.5% to 24% (Figure 3F; p<0.01). Interestingly, both species exhibited a shift towards longer polysomes (Figure S6D).

In summary, the overall non-significant change in ribosome concentrations, and the opposing trends of changes to the polysome fractions observed for the two yeast species at 1h ED do not reflect consistent global changes in molecular crowding. We conclude that while molecular crowding may exert local effects, it is unlikely to explain global cytosolic solidification.

### Cytosolic acidification triggers FAS assembly

Energy depletion is known to lead to a drop in intracellular pH in yeast^65^, which correlates with reduced particle mobility and the widespread assembly of cytoplasmic proteins into foci^26^. We thus attempted to probe the effect of acidification on foci assembly, and disentangle it from a potential effect of intracellular crowding induced by a reduction in cellular volume.

We first used live-cell confocal microscopy to quantify cell volumes under the following conditions: native state, ED, osmotic stress (hyperosmolar condition: 1.2 M sorbitol), and ectopic acidification (addition of 2 mM 2,4-dinitrophenole (DNP)) in the naturally acidic growth medium (pH 4.5) (Figure 4A, Methods). The volumes measured for native state cells were in line with the 80-100 µm^3^ reported for *S. pombe*^106^ and the roughly 40 µm^3^ reported for *S. cerevisiae*^107^. There were no significant changes to cell volumes upon ED, in line with our tomography-based findings on marginal changes in intracellular ribosome concentrations.

**Figure 4.**
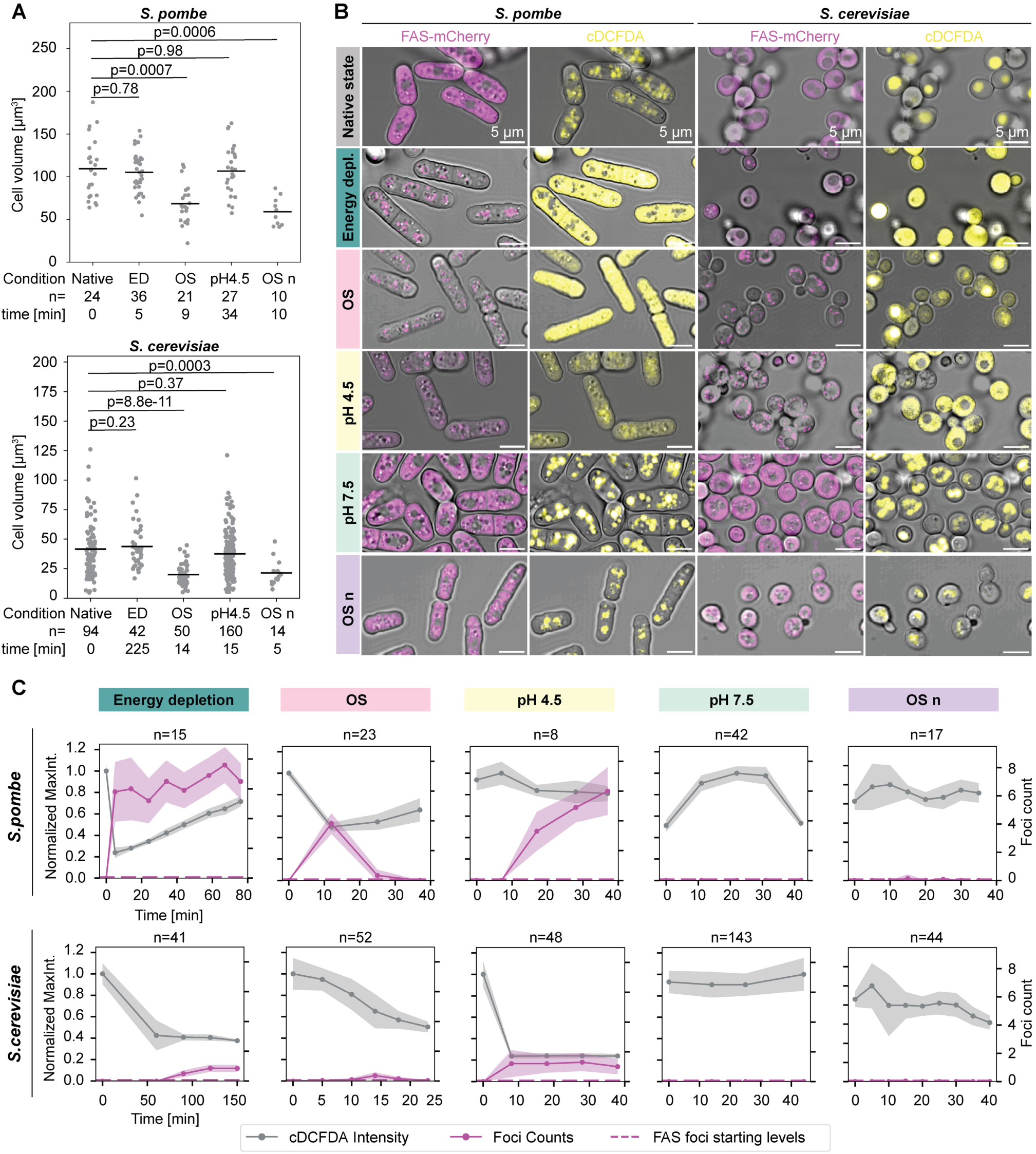
Acidification drives FAS assembly under different stress conditions. (A, B) Brightfield and confocal microscopy stacks of cells tagged with FAS-mCherry and co-stained with the acidic pH indicator CDCFDA acquired in native state and under ED, osmotic stress (OS), ectopic acidification (pH <4.5) and osmotic stress in pH 7.5 buffered medium (OS n). (A) Quantification of cell volumes under the different conditions reveal a significant decrease only under hyperosmolar conditions. Statistical significance (p<0.05) performed with Kolmogorov–Smirnov test. Cell numbers (n) are indicated for each condition. (B) Central confocal slices showing FAS-mCherry and CDCFDA channels overlaid on brightfield images. (C) Quantification of FAS foci count per cell (magenta, right y-axis) plotted alongside corresponding maximum intensity of the CDCFDA signal (grey, left y-axis) over time under different stress conditions. Data plotted as average (solid lines) ± SD (shaded area). The average number of cells analysed for each timepoint are indicated (n). Dotted lines: starting levels of FAS foci counts. Overall, more foci formed in *S. pombe* compared to *S. cerevisiae* under the same conditions.

As expected, upon hyperosmotic stress, the volumes of *S. pombe* and *S. cerevisiae* cells were significantly reduced by 38% and 52% in the natural acidic cell medium. Similarly, hyperosmotic stress performed under growth medium buffered to neutral pH showed significant cell size reduction. Cell volumes of ectopically acidified yeast were similar to those in native state.

Next, we monitored FAS assembly dynamics in relation to intracellular pH under the different conditions by co-staining with carboxy-DCFDA as an indicator for acidic environments (Figure 4B, C, Figure S7). In native state, FAS was dispersed throughout the cytosol and CDCFDA stained only the acidic vacuoles. Upon ED, FAS assembled into foci, coinciding with acidification of the cytosol (Figure 4B). As reported above, FAS condensed in *S. pombe* immediately after stress induction, while foci formation was delayed in *S. cerevisiae* (Figure 2C, 4C). Under osmotic stress, FAS foci formed within 10 minutes and dissolved within a similar time frame (Figure 4B, C), suggesting an adaptation to the stress. Surprisingly however, osmotic stress also induced cytosolic acidification that coincided with FAS assembly into foci. While this was observed for all *S. pombe* cells, only a subset of the *S. cerevisiae* cells reacted to this treatment. Under ectopic acidification using the natural acidic cell medium, FAS assembled within 10 minutes in both species. The volume of these cells was similar to that in the control (Figure 4A), suggesting no large-scale changes in macromolecular crowding that could trigger foci formation. To validate that the pH drop is indeed the driver for FAS assembly, we repeated the experiment (addition of DNP) with the growth medium buffered to pH 7.5 or 5. At neutral pH, FAS remained dispersed in the cytosol of both yeast species similar to the native state (Figure 4B, C; Figure S7A, B), whereas foci formed at pH 5 (Figure S7A-C). To fully disentangle the effects of cytosolic acidification and crowding, we tested whether FAS condenses into foci in response to hyperosmolar conditions when the surrounding medium is buffered to pH 7.5 (Figure 4B). While the osmotic stress led to an expected decrease in cell volume of 54% and 51% in *S. pombe* and *S. cerevisiae*, respectively (Figure 4A), FAS did not form foci at neutral pH. Thus, we conclude that acidification is the main driver of FAS assembly in different stress conditions.

The isoelectric point (pI) of FAS, *i.e.* the pH at which the complex exhibits minimum solubility due to minimal electrostatic repulsion, is in the acidic range (*S. pombe*/*S. cerevisiae* FAS1: 6.0^108^/5.7^109^, FAS2: 5.8^110^/5.1^111^; theoretical pIs for FAS1/2 of *S. pombe were* calculated based on UniProt sequences Q9UUG0^108^/Q10289^110^ with the ExPASy Compute pI/Mw^112^). Consequently, in the acidified cytosol under different stress conditions, the pH is closer to the pI of FAS, potentially promoting the condensation of the complex either through self-assembly or interactions with additional partners. This is in agreement with FAS surface charges exhibiting more neutral and positive patches at pH 4.5 compared to mildly negative patches at pH 7.5 (Figure S7E, F).

To conclude, we show that different types of exogenous stress induce a pH drop in both of the evolutionary-distant yeast species and that induction of controlled acidification is sufficient to trigger mesoscale assembly as probed here by light microscopy of FAS, irrespective of changes in cell volume. These results support previous reports indicating acidification to be the trigger for cytosolic solidification^26^.

### Supramolecular assembly as part of the general yeast stress response

To investigate whether the osmotic stress (in naturally acidic growth medium) and ectopic acidification conditions that triggered FAS foci also lead to broader structural reorganization of the yeast cytosol, we acquired cryo-ET data of the cells under these conditions (Figure 5, Table 1).

**Figure 5.**
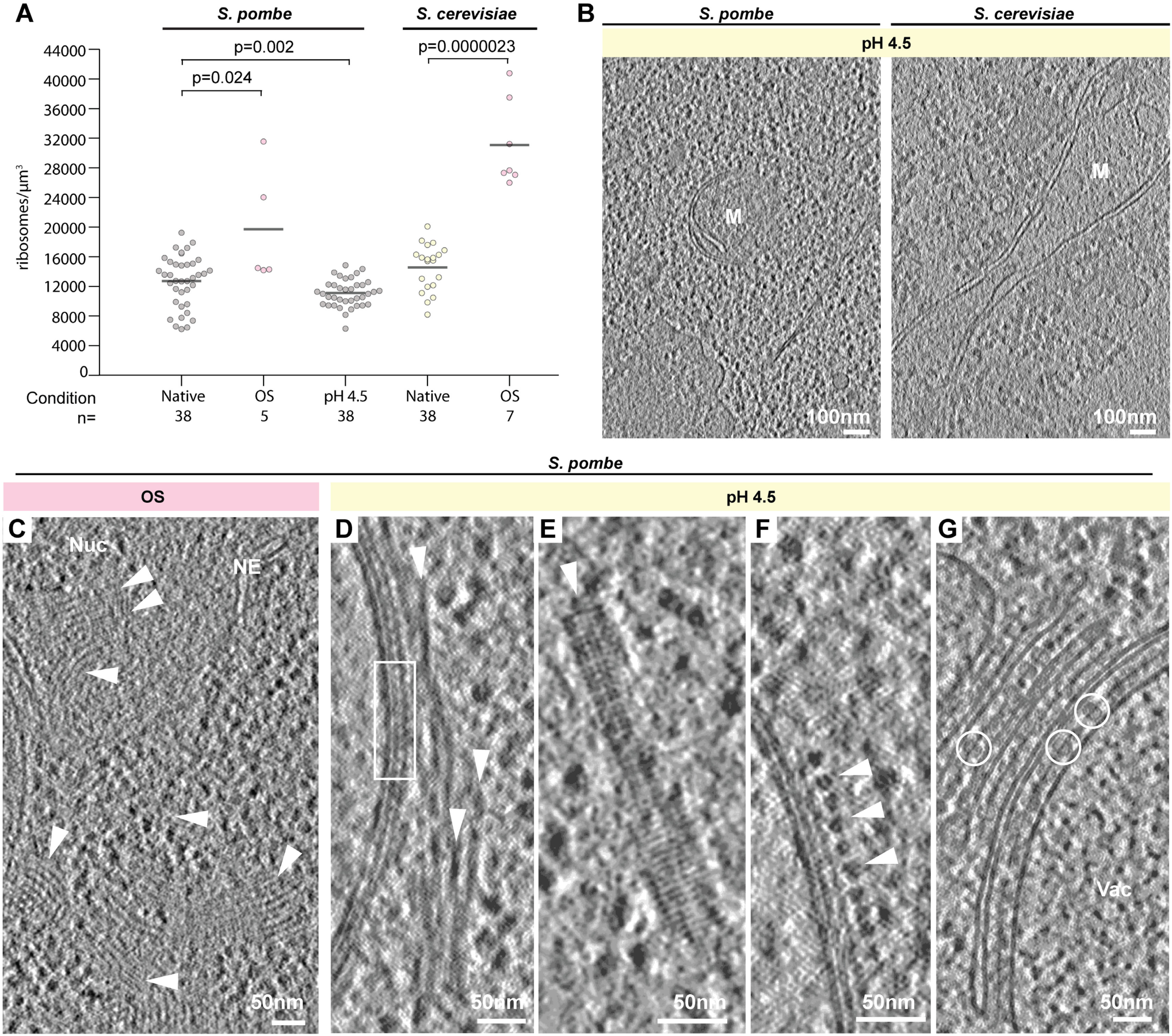
Supramolecular assemblies form upon cytosolic acidification. (A) Ribosome concentrations determined based on particle numbers per cytosolic volume in tomograms of osmotically stressed and ectopically acidified cells compared to respective native state (native state data replotted from Figure 3B). Statistical significance (p<0.05) performed with Kolmogorov–Smirnov test. Tomogram numbers (n) are indicated for each condition. (B) Ribosomes are discernible in acidified *S. pombe*, while ectopic acidification of *S. cerevisiae* leads to aggregation of the cytoplasm. M = mitochondrion. (C) Lattice-like nuclear and cytosolic structures (white arrowheads) observed in tomograms of *S. pombe* cells subjected to hyperosmolar conditions for 10 minutes. NE: nuclear envelope. (D-G) Supramolecular assemblies (white arrowheads) observed in response to ectopic acidification in *S. pombe*. A surface lattice connected to the outer mitochondrial membrane is indicated by a white box in D. (G) FAS (white circles) forms monolayer arrays between membrane stacks. Vac = vacuole.

First, we quantified the ribosome concentrations to examine and validate changes in intracellular crowding at the subcellular level (Figure 5A). In line with our cell volume measurements (Figure 4A), ribosome concentrations significantly increased under osmotic stress, with an average of 19,704 ± 7,847 ribosomes/µm^3^ and 31,075 ± 5,818 ribosomes/µm^3^ for *S. pombe* and *S. cerevisiae.* These concentrations correspond to a 1.6 and 2.1-fold increase compared to the native state, in agreement with cell size reduction of 38% and 52% for *S. pombe* and *S. cerevisiae,* respectively. These data illustrate that measurements of ribosome concentrations from cryo-ET data accurately reflect intracellular crowding levels. Upon ectopic acidification, the average ribosome concentration in *S. pombe* cells was 10,947 ± 1,769 ribosomes/µm^3^, thus reflecting a 12.5% decrease in intracellular crowding compared to the native state (12,708 ± 3,460 ribosomes/µm^3^), similar to the slight decrease observed in 1h ED. Of note, the cryo-ET data of ectopically acidified yeast revealed striking differences between the ability of the two species to cope with low cytosolic pH. The cytosol of *S. pombe* was populated with discernible ribosomes and ribosome-excluded areas after 50 minutes of acidification (Figure 5B, Figure S7D). In contrast, the cytosol of *S. cerevisiae* appeared aggregated within 10 minutes of acidification, precluding quantification of ribosome concentrations (Figure 5B, Figure S7D). This unexpected observation suggests that more physiological stressors, including starvation and osmotic stress, despite leading to cytosolic acidification, elicit mechanisms that allow adaptation and survival that are different from those in ectopic acidification.

We next examined the presence of mesoscale assemblies, such as those observed in ED (Figure 1E-K), under osmotic stress and ectopic acidification. Under hyperosmotic conditions, *S. pombe* exhibited polymeric assemblies within 5-15 minutes (Figure 5C), the timeframe within which FAS foci and cytosolic acidification were observed by confocal microscopy (Figure 4B, C).

The polymeric assembly was present in both cytosol and nucleus, and resembled the ED polymer observed in *S. cerevisiae* (Figure 1I). Overall, 18% of *S. pombe* tomograms contained filamentous structures in osmotically stressed cells (Table 1), while FAS assemblies were not observed, likely due to their transient nature (Figure 4B). We did not observe higher-order assemblies in the tomograms of osmotically stressed *S. cerevisiae*, possibly due to the lower number of cells that reacted to this stress condition as shown by light microscopy (Figure 4B, C).

In tomograms of ectopically acidified *S. pombe*, we identified three different polymeric assemblies (Figure 5D-F), including the dynamin-like lattices on the surface of mitochondria (Figure 5B, F). One of the remaining two exhibited lattice-like organization (Figure 5E) of 4 nm subunits and resembled an architecture observed in both yeast species under ED (Figure 1H). Finally, FAS complexes organized into 2D monolayers staked between membranes (Figure 5G), identical to their organization in ED *S. pombe* (Figure 1J). The aggregation of the acidified *S. cerevisiae* cytosol (Figure 5B) precluded the detection of supramolecular assemblies with defined morphologies.

In summary, structural reorganization of the cytosol by supramolecular assembly, as we have characterized for ED, occurs also under osmotic stress and ectopic acidification. Changes to ribosome concentration that report on global changes to intracellular crowding do not appear to be a common response across the different stress conditions. Our findings thus pinpoint acidification as a common effect of all examined stress conditions leading to structural reorganization and supramolecular assembly in yeast.

## DISCUSSION

Here, we investigated the basis of the phenomenon of cytoprotective solidification reported in yeast stress response using label-free cryo-ET of cells minimally perturbed by specimen preparation. In agreement with previous studies, we confirmed that changes to organelle morphologies^73,74,106,113^ and formation of higher-order molecular assemblies occur under ED^26,27^. Previous genome-wide screens in *S. cerevisiae* identified over 200 proteins to assemble into foci following nutritional stress, most of which are either metabolic enzymes, RNA binding proteins or translation factors^35,40^. Whether such foci represent dysfunctional protein aggregates or storage compartments that segregate active or inactive enzymes, remained unknown. While our molecular-resolution data revealed the presence of only a handful of structured assemblies, our analyses provide insight into these aspects of supramolecular assembly in response to stress.

Our cryo-electron tomograms of *S. pombe* and *S. cerevisiae* cells under ED, OS and ectopic acidification showed both common and species-specific structured assemblies, most of which remain of unknown identity. Potential candidates include translation factors, such as eIF2B^37^, and essential metabolic enzymes, such as glutamine synthetase or CTP synthetase^38^. Emerging combinations of whole-cell mass spectrometry-based proteomics^114–119^ and multimeric structure prediction^120–122^ may be used in the future for comparison with higher-resolution density maps generated from in-cell cryo-ET^123^ to help unambiguously identify assembly forming proteins. A recent example of such approaches includes the FilamentID method^45^. We further noted the presence of unstructured assemblies, which could only be inferred from the exclusion of ribosomes. As these are likely formed by more flexible and possibly disordered macromolecules^31,124–129^, cryo-ET can only provide limited structural information. An interesting exception were assemblies of FAS complexes that could be identified based on their stereotypical structural signature. In *S. pombe,* FAS arranged into ordered monolayer arrays staked between membranous compartments under ED and acidification. This array organization is likely induced by membrane association, through yet unidentified interaction partners that associate with the complex under stress conditions. In contrast, the complex organized into clusters of seemingly random orientations in ED *S. cerevisiae*. Our structural analysis confirmed that the condensed complex is fully assembled in both architectures under ED, and the positioning of densities corresponding to the acyl carrier proteins at the top of each half dome supports a catalytically inactive state of an otherwise intact enzyme^86^. Hence, FAS mesoscale assembly correlates with inactivation of the enzyme upon ED. This is in line with the notion that enzymes are sequestered into assemblies as functional units that can resume their catalytic activity when cellular stress subsides^27^. It remains to be determined whether inactivation of FAS is related to low levels of metabolites and/or substrates under ED, or whether other regulatory mechanisms may be at play.

Similarly, ribosomes at 1h ED in both yeast species comprised assembled 80S complexes with predominantly empty PTCs, indicating a non-translating state. It is somewhat surprising that ribosomes are maintained as fully assembled 80S complexes in starved and ED cells, as non-translating ribosomes are expected to disassemble into free large and small subunits, while the released mRNA is sequestered into stress granules^130^. This indicates specific mechanisms for ribosome hibernation under the examined stress conditions. The moderate resolutions of the ribosome maps obtained in our work however precluded identification of densities that may explain the stabilization of the two ribosomal subunits in the seemingly empty 80S complex. Nevertheless, our finding is in line with recent cryo-EM studies of cytosolic ribosomes isolated from yeast showing vacant PTCs. In glucose-depleted *S. pombe*, a conformational change in the PTC is reported to disrupts the inter-subunit bridge B2a, potentially occluding tRNAs and mRNA from binding, and thereby inhibiting translation^75^. In starved *S. cerevisiae*, the protein Stm1 at the inter-subunit interface is shown to clamp the two subunits together, preventing their dissociation and simultaneously inhibiting translation by exclusion of mRNA binding^131^. We hypothesize that preserving fully assembled but inactive ribosomes under critically low energy conditions, similarly to the FAS complexes discussed above, could represent a protective storage mechanism. This would allow for energy-efficient re-entry once favorable physiological conditions are restored.

We further aimed to untangle the physicochemical factors that underpin yeast cell solidification. Macromolecular crowding and acidification have been proposed as potential triggers for this phase transition, that is experimentally manifested in reduced molecular mobility^25,26,59^. We evaluated changes in crowding by quantifying ribosome concentrations and polysome fractions, representing the most prominent molecular species implicated in intracellular crowding. Ribosome concentrations at 1h ED were similar to native state cells. Previous polysome profiling experiments in *S. cerevisiae* show that polysomes disassemble within 10 minutes of energy depletion^37^ or glucose deprivation^47,61^. This initial polysome collapse contributes to an increase in mesoscale particle mobility, and is shown to be the main factor leading to transient fluidization of the cytoplasm^47^. Beyond 15 minutes of starvation, particle diffusivity gradually decreases^47^, ultimately reproducing the previously described solidification in both *S. cerevisiae* and *S. pombe*^25,26^. After 60 minutes of starvation, polysomes reassemble in *S. cerevisiae* to restart translation of stress response-related proteins^61^. Our cryo-ET data showed that polysome fractions decreased at 1h ED in *S. pombe* with most ribosomes showing empty PTCs, but increased in *S. cerevisiae* where 20% of the ribosomes contained P-site tRNAs indicative of active translation. Taken together, our findings of minor changes in ribosome concentrations and the opposing trends in polysome fractions across the two yeast species do not support the suggested mechanism of a uniform increase in molecular crowding as the basis for solidification in yeast cells upon nutrient deprivation.

We thus additionally investigated the impact of acidification. Common cellular stresses, such as nutrient restriction and heat shock, coincide with cytosolic acidification^25,26,67^. Acidic conditions lead to condensation of essential RNA-binding proteins involved in translation regulation, like Pab1, Ded1 and Pub1, and their sequestration into stress granules^31,132,133^. Such phase transitions are important in promoting survival under stress. Acidification further increases resistance to stress in *S. cerevisiae*, acting as a prerequisite for activation of heat shock proteins^67^. Following on work by Munder et al.^26^, we show that artificially-induced cytosolic acidification, even in the presence of an energy source, was sufficient to induce the condensation of FAS, used here as a model for essential metabolic enzymes, and the formation of various structured mesoscale assemblies. At the same time, cell volumes and ribosome concentrations remained similar to native state cells. We further showed that osmotic stress, in addition to its expected effect of significantly reduced cell volume and increased intracellular crowding, also resulted in a sharp pH drop that coincided with FAS foci formation. Importantly, FAS did not condense under hyperosmolar conditions when the cells interior was maintained at neutral pH. Thus, cytoplasmic acidification appears to be a general response to vastly different stresses in yeast, and we show that the mere acidification of the cytosol induces the formation of a number of structured assemblies.

At lower intracellular pH, the surface charge of proteins with isoelectric points in the acidic range is reduced. As a consequence, surfaces become less solvated, increasing the probability of self-association or the formation of new interactions^55^. In fact, the isoelectric points of a large fraction of the yeast cell proteome clusters in the acidic range^26,67^, suggesting that many proteins become less soluble at low pH. Similar to what has been reported for *E. coli*, yeast proteomes may have also evolved at the edge of metastability in terms of their propensity to self-associate^134^. This enables cells to exploit physicochemical cues such as acidification to promote (self) assembly, and thereby to establish a protective mechanism for stress survival^37^.

Self-assembly of proteins into mesoscale structures across the cytoplasm can collectively contribute to an overall increase in crowding and to changes in the cells’ physical properties^25,26,135^. The higher-order structured assemblies we observed, while likely reflecting only a fraction of the cellular proteome that undergoes assembly and condensation under stress, occupy a relatively large fraction of the cytosolic volume (around 5% in 1h ED cells). An increase in the volume fraction of larger ordered structures, and likely of disordered condensates, can impact the mesoscopic properties of the cytosolic fluid on different length scales. On the one hand, molecular assembly can lead to jamming of the constituent particles that become kinetically trapped^136,137^. On the other hand, mesoscale structures can have an effect on the connectivity of the fluid phase, leading to confinement and constrained diffusion of non-interacting molecules. To determine what scale of molecular assembly and volume exclusion is sufficient to explain the significantly reduced particle diffusion reported under stress conditions will require modeling at multiple spatial and temporal scales^46,138^. Our work provides structural, quantitative and spatial data that can contribute towards the generation of cross-scales models^139^, to enable a holistic understanding of cellular organization and the effect of perturbations from the molecular to the whole-cell level.

## ACKNOWLEDGMENTS

We thank the EMBL cryo-EM platform, in particular Wim Hagen and Felix Weis, Thomas Hoffmann and EMBL IT for computational support, the EMBL Advanced Light Microscopy Facility, especially Manuel Gunkel and Aliaksandr Halavatyi, Frosina Stojanovska for her support with DeePiCt, Christian Haering and Axel Mogk for their kind gift of cells and plasmids, Christian Zimmerli, Matteo Allegretti, and Martin Beck for kindly sharing *S. cerevisiae* cryo-ET data, Tobias Walther for revising template matching results for ribosomes and Alexander Mattausch for revising organelle segmentations. We thank Simon Alberti, Titus Franzmann and Liam Holt for critical reading of the manuscript and valuable discussion. M-C.S was supported by a postdoctoral fellowship from the National Science Foundation Center for Quantitative Cell Biology. R.K.J. was supported by a postdoctoral fellowship from the Independent Research Fund Denmark (0106-00010A). This work was supported by the EMBL, a European Research Council Starting Grant (3DCellPhase^-^ 760067), Chan Zuckerberg Initiative grants for visual proteomics (2021-234620, 2023-331612), and the National Science Foundation Science and Technology Center for Quantitative Cell Biology (NSF-STC grant 2243257).

## AUTHOR CONTRIBUTIONS

J.M. conceived this study. S.K.G. and M-C.S. collected and analysed the light-microscopy and cryo-ET data. E.Z. created the *S. pombe* FAS-mCherry strain and assisted in confocal microscopy data collection, experiment design and FIB milling. R.K.J. supported the subtomogram averaging. W.S. assisted in light-microscopy data acquisition and analysis. M-C.S., S.K.G., and J.M. wrote the manuscript with input from all co-authors.

## DECLARATION OF INTERESTS

The authors declare no competing interests.

## STAR METHODS

### Key resources table

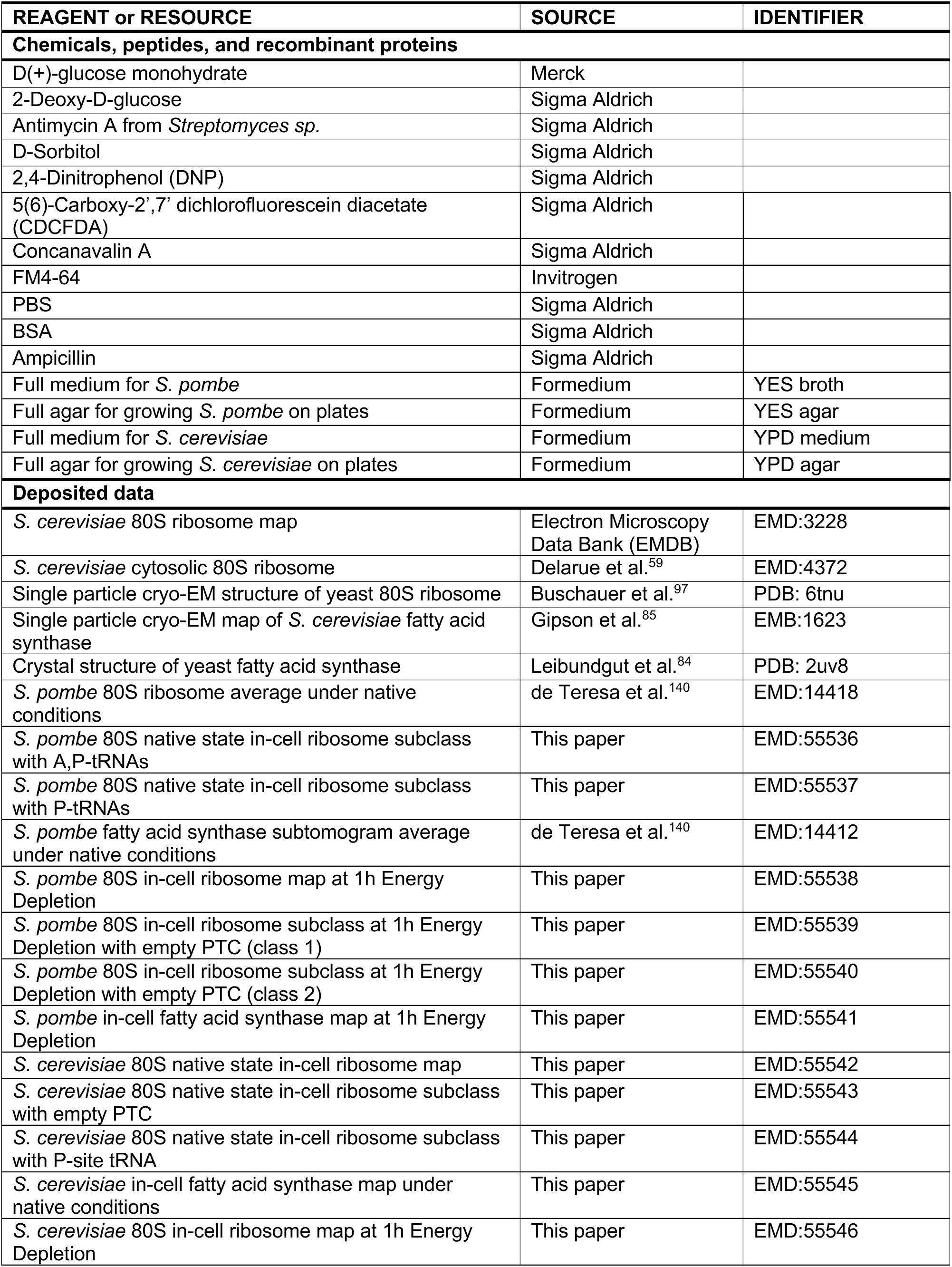
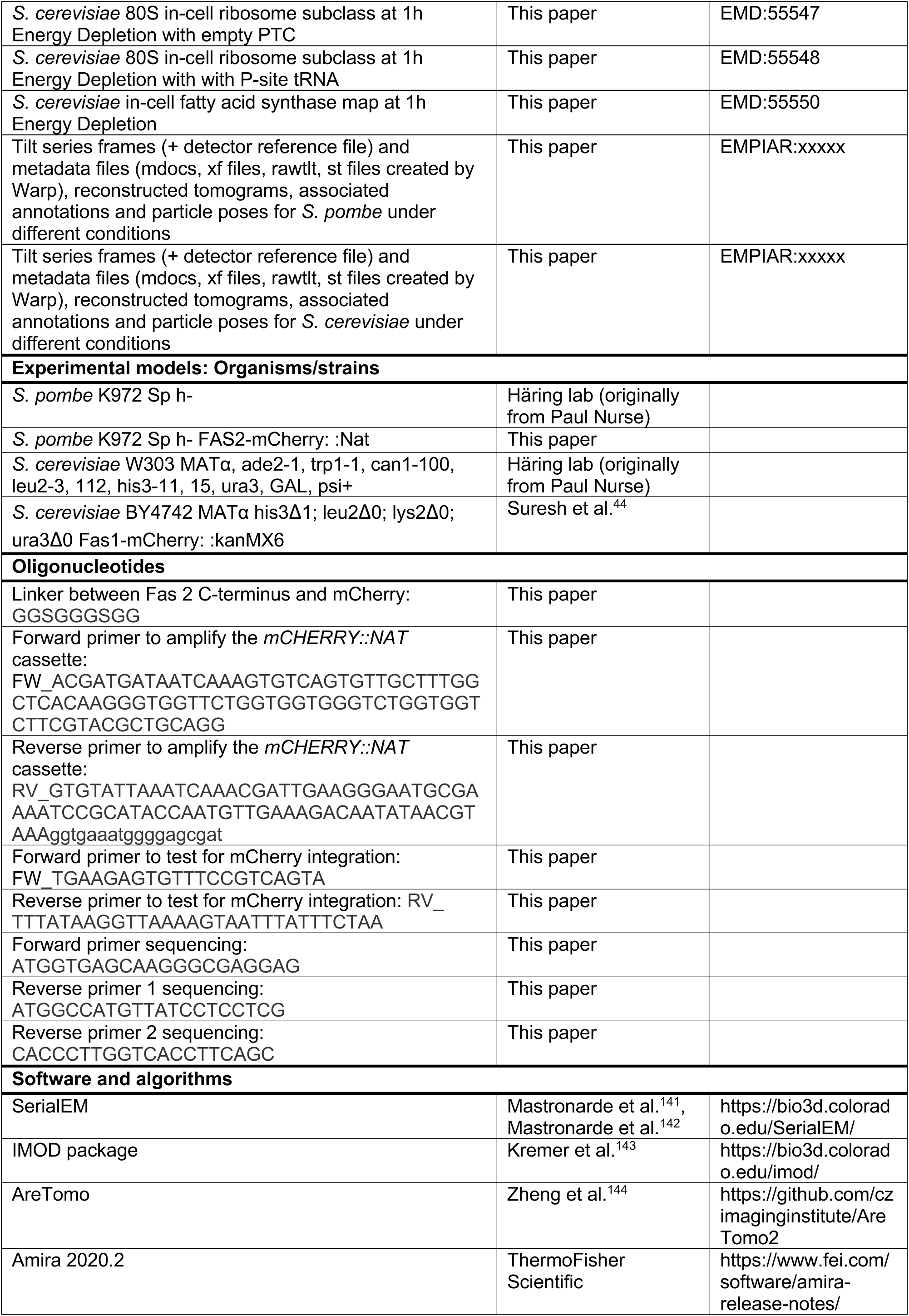
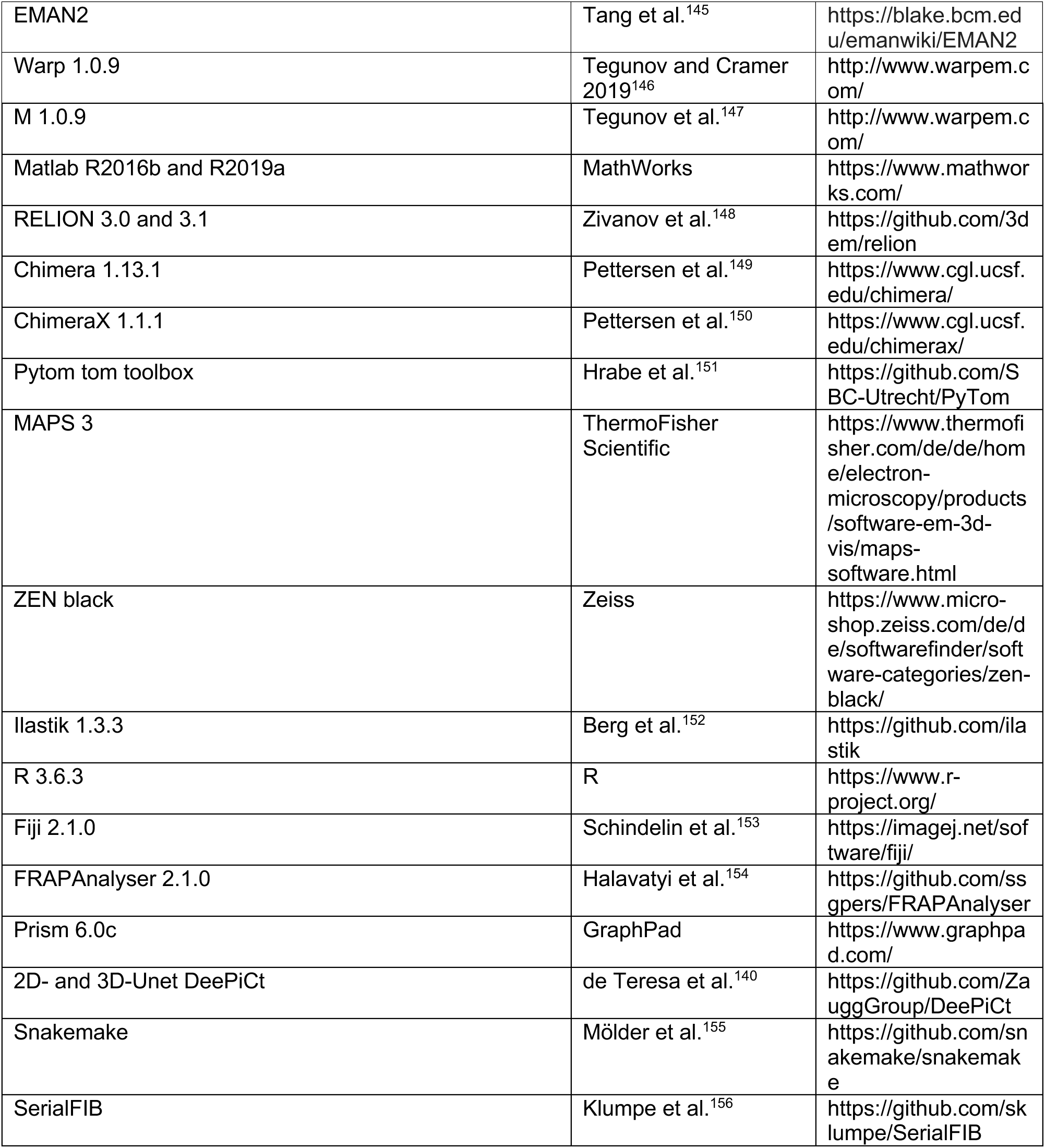

## RESOURCE AVAILABILITY

### Lead contact

Further information and requests for resources and reagents should be directed to and will be fulfilled by the lead contact, Julia Mahamid.

### Materials availability

All unique materials and reagents generated in this study are available from the lead contact with a completed material transfer agreement.

### Data and code availability

- Atomic models of the *S. cerevisiae* FAS and ribosome were obtained from the Protein Data Bank (PDB): 2uv8 and 6tnu, respectively, and density maps from the Electron Microscopy Data Bank (EMDB): EMBD:1623 (*S. cerevisiae* FAS), EMD:3228 and EMD:4372 (*S. cerevisiae* ribosome).
- All cryo-ET data generated and analyzed in this study will be deposited in the Electron Microscopy Public Image Archive. Subtomogram averages of ribosomes and FAS are deposited in the EMDB. All data will be released upon acceptance of this manuscript, with accession numbers provided in the key resource table.
- This paper does not report original code.
- Any additional information required to reanalyze the data reported in this paper is available from the lead contact upon request.

### Experimental model

*Schizosaccharomyces pombe* and *Saccharomyces cerevisiae* strain backgrounds were K972 (h-), W303 (ade2-1, trp1-1, can1-100, leu2-3,112, his3-11,15, ura3, GAL, psi+) or BY4742 (his3Δ1; leu2Δ0; lys2Δ0; ura3Δ0). The *S. pombe* Fas2-mCherry expressing strain was constructed via homologous recombination-directed PCR-based one-step C-terminal tagging^44^. Between the Fas2 C-terminus and mCherry, a linker of the following sequence was introduced: GGSGGGSGG. Primers used to amplify the *mCHERRY::NAT* cassette were as follows: FW_ACGATGATAATCAAAGTGTCAGTGTTGCTTTGGCTCACAAGGGTGGTTCTGGTGGTGGGTCTGGTGGTCTTCGTACGCTGCAGG, RV_GTGTATTAAATCAAACGATTGAAGGGAATGCGAAAATCCGCATACCAATGTTGAAAGACAATATAACGTAAAggtgaaatggggagcgat.

CloNat resistant clones were tested for correct mCherry integration by PCR with the following primers: FW_TGAAGAGTGTTTCCGTCAGTA and RV_TTTATAAGGTTAAAAGTAATTTATTTCTAA, and the genomic DNA of the respective clones used as a PCR template. The PCR product was sequenced with the following primers: ATGGCCATGTTATCCTCCTCG (mCherry RV-1; anneals at N-terminus), CACCCTTGGTCACCTTCAGC (mCherry RV-2; anneals close to the middle of the mCherry coding sequence) and ATGGTGAGCAAGGGCGAGGAG (mCherry FW; anneals at the N-terminus of the mCherry coding sequence).

## METHOD DETAILS

### Yeast strains and media

*S. pombe* and *S. cerevisiae* were recovered from frozen stocks (stored at -80°C) and streaked onto YES or YPD agar plates, respectively. Cells were either incubated on the agar plates at 30°C for 1-3 days or kept at room temperature for three days. Cells were re-streaked onto fresh plates from single colonies and incubated for 1-2 days as described above. Single colonies were then picked from these plates to inoculate 5-20 ml of complete media (YES or YPD, respectively) or species-tuned synthetic media supplemented with 20 mM dextrose. The liquid cultures were incubated at 30°C overnight with shaking at 165 rpm (NCU-Shaker mini, Benchmark) in glass Erlenmeyer flasks that fit approximately 10 times the volume of the culture. Cells were grown to the exponential phase and used in experiments at an OD600 between 0.2 and 0.6.

### Stress treatment

Yeast cells were harvested during their exponential phase of growth. Native state cells (control) were washed thrice in synthetic medium (EMM without dextrose for *S. pombe* and complete supplement mixture plus yeast nitrogen base for *S. cerevisiae*) supplemented with 20 mM Glucose. For energy depletion, cells were washed thrice in synthetic media without dextrose containing 20 mM 2-deoxyglucose and 10 mM Antimycin A to simultaneously inhibit glycolysis and mitochondrial respriation^23,26,27^. To induce osmotic stress, cells were washed thrice in synthetic medium supplemented with 20 mM dextrose and 1.2 M sorbitol^26,135^. Where indicated, the hyperosmolar medium was additionally buffered to pH 7.5. For experiments with pH adjusted conditions, cells were washed thrice in synthetic medium with 20 mM dextrose adjusted to pH 5 or pH 7.5^26,29^. After the final washing step, 2 mM 2,4-dinitrophenol was added to the pH adjusted media to equilibrate the intra-and extracellular pH. Each washing step consisted of centrifugation at 4,000 rpm for 3 minutes (Eppendorf 5424R centrifuge), discarding of the supernatant and replacing it with 1 ml of the respective medium.

### Plunge freezing for cryo-ET

Plunge freezing was carried out using a Leica EM GP2 (Leica Microsystems). Grids were glow discharged immediately before plunging on both sides for 45 s at 0.26 mbar and 15 mA (Pelco Easy glow). 4 µl of yeast cell suspension at an OD600 of 0.3 (energy depletion) or 0.6 (osmotic stress and different pH conditions) were deposited on quantifoil holey carbon or silicon dioxide support film side of the copper grids (Quantifoil R2/1, Cu 200 mesh grid, Quantifoil Micro Tools). Excess liquid was removed by blotting from the back of the support film side for 1-2 seconds at 22°C and a humidity of 99%. Grids were plunge-frozen into liquid ethane cooled to -183 to -185°C, and stored in liquid nitrogen until further use.

### Cryo-FIB milling

Plunge-frozen grids were mounted under liquid nitrogen into custom AutoGrids with a cutout allowing milling at shallow ion beam incident angles. Grids were then placed inside a 45° pre-tilt shuttle and transferred into a dual-beam cryo-FIB-SEM microscope (Aquilos, Thermo Fischer Scientific). An initial sputter-coating with platinum for 10-20 s (1 kV, 10 mA, 10 Pa) and a coating with organometallic platinum for 8-11 s at a stage height of 10.6-11.6 mm using the gas injection system were applied to render the grids conductive and reduce curtaining artifacts during subsequent ablation of cellular material, respectively. Suitable positions for FIB-milling of lamellae were determined manually and eucentricity was refined in the MAPS software (Thermo Fisher Scientific). Lamellae were prepared in consecutive steps of rough and fine milling at a stage angle of 15°-20°, resulting in lamellae with 8-13° pretilt, and a constant ion beam voltage of 30 kV. Rough-milling was carried out either manually or using the SerialFIB software^156^ in three steps to create subsequently thinner lamellae of 5 µm, 3 µm and 1 µm using 1 nA, 0.5 nA and 0.3 nA currents, respectively. The milling process was monitored with SEM imaging (10 kV, 50 pA). Rough-milled lamellae were manually thinned down to the target thickness of around 200 nm with the focused ion beam at 50 pA. Grids were sputter coated with platinum for 3-5 s (1 kV, 10 mA, 10 Pa), removed from the microscope and stored in liquid nitrogen until cryo-electron tomography acquisition.

### Cryo-ET

Cryo-ET data were acquired on a Titan Krios microscope (Thermo Fischer Scientific) operated at 300 kV, and equipped with a field emission gun, a Quantum post-column energy filter (Gatan) and either a K2 Summit or a K3 direct electron detector (Gatan). Data were collected in zero-energy loss and dose-fractionation modes. Tilt series were acquired using automation scripts in SerialEM^141,142^ at a detector-dependent magnification of 42,000x or 26,000x (calibrated pixel size depending on the detector: 3.37 or 3.43 Å, respectively) and 2-4 µm defocus from -60° to +60° in 2° increment following a dose-symmetric tilt scheme^157^ starting from the lamella pre-tilt. The total accumulated dose on the sample was kept below 150 e^-^/A^2^. Defocus-only tomograms were collected with a 70 µm objective aperture. For a subset of the data, a Volta phase plate^158^ (VPP, Thermo Fisher Scientific) was used after conditioning for around 5 minutes.

The data analysed in this work is detailed in Table 1. In total, 48 *S. pombe* and 64 *S. cerevisiae* native state tomograms were acquired. For 1h ED, 117 *S. pombe* and 62 *S. cerevisiae* tomograms were acquired. For the later timepoints of ED, the following numbers of tomograms were acquired: 76 tomograms for 3.5h ED and 34 tomograms for 17h ED of *S. pombe*, and 56 tomograms for 6h ED of *S. cerevisiae*, For *S. pombe*, additional 31 tomograms were acquired after 4 days of glucose depletion. For the osmotic stress, we acquired 43 tomograms for *S. pombe* at 15 minutes and 22 tomograms for *S. cerevisiae* at 10 minutes. For the ectopic acidification condition (equilibration of intracellular pH to that of acidic synthetic medium supplemented with 20 mM glucose), we acquired 10 tomograms of *S. pombe* after 8 minutes and 66 tomograms after 50 minutes of acidification. Of the latter tomograms, 27 were acquired at a different pixel size from the rest of the data set (2.075 Å). These data were used for visual inspection of supramolecular assemblies, but excluded from the ribosome concentration quantification. For *S. cerevisiae*, we acquired 13 tomograms after 35 minutes of acidification, all exhibiting major aggregation that precluded further analysis.

### Tomogram reconstruction

Prior to tilt-series alignment, tilt movie frames were corrected for detector background signal and beam-induced motion in Warp 1.0.9^146^. The pre-processed tilts were exported for alignment and reconstruction in IMOD^143^ either by patch tracking or using platinum speckles deposited during the sputtering as fiducials. Tomograms were reconstructed at 4x binning in IMOD^143^ or AreTomo^144^ corresponding to a pixel size of 13.48 Å or 13.7 Å, by weighted back-projection, for segmentation and particle localization.

### Tomogram segmentation

Segmentations of cellular structures were conducted with DeePiCt^140^, which employs a 2D and a 3D Convolutional Neuronal Network (CNN). The 2D CNN was used for supervised segmentations of the different organelles, ED assemblies and the cytosolic volume. The 3D CNN was employed to localize macromolecules, such as ribosomes and FAS complexes and to predict membranes (described further below). For training of the 2D CNN, ground truth annotations of cellular features and boundaries of the cytosol were generated by manual segmentation in a total of 20 tomograms (ten tomograms acquired using the VPP with defocus, and ten defocus-only tomograms) in Amira (Thermo Fisher Scientific) as previously described^140^. 3D segmentations were interpolated from annotations on every 5^th^ to 15^th^ 2D tomographic slice and cleaned manually. Both the 2D and 3D trained models are available on the DeePiCt^140^ Github repository (https://github.com/ZauggGroup/DeePiCt#Models). Automated organelle and cytosol detection were performed using the DeePiCt^140^ 2D CNN (D=5, IF=16) on 4x binned tomograms that were pre-processed by amplitude spectrum matching (low-pass cut-off at 350 pixel, sigmoid curve smoothing cut-off of 20 voxel in Fourier space). Predictions were generated in patches of 288 x 288 voxel cropped to 40 voxel and with a z-cut-off of 200 slices (100 slices above and below the z-center). The prediction outputs were assembled into 3D volumes from 2D tiles, filtered with a 1D Gaussian along the z-axis with a sigma of 5 and evaluated with an output threshold score of 0.7.

### Volume occupancy of cytoplasmic assemblies

The volume occupancy of structured assemblies in tomograms of 1h ED cells was determined by dividing the segmented volumes of the structures by that of the cytosol (described above). Voxel-based volumes were calculated by integrating segmented voxels in the binary masks using MATLAB and converted into physical units of µm^3^ by multiplication with pixel sizes and binning factors.

### Ribosome localization, subtomogram averaging and distribution analysis

For subtomogram averaging, 80S ribosomes were localized through a combination of template matching using PyTom^151^, DeePiCt^140^-based 3D CNN predictions and manual annotation acquired on defocus-only tomograms (described in detail in Trueba et al.^140^). Specifically, ribosomes were comprehensively localized with this approach in 10 *S. pombe* and 9 *S. cerevisiae* native state tomograms, and in 20 *S. pombe* and 16 *S. cerevisiae* 1h ED tomograms. Tomograms were first filtered by amplitude spectrum matching to generate high image contrast prior to running predictions with a pre-trained ribosome models (depth=2, initial features=4, batch normalization) with a box size of 64 voxels. A threshold cut-off at 0.5 of the output score was applied to the prediction results. The centroids were determined from clusters with a minimal size of 100 voxels. False positives outside the cytosolic volume were removed by multiplication with a cytosol mask considering a contact size of 10 pixels. The cytosol masks were either generated manually through segmentation in Amira (Thermo Fisher Scientific) or resulted from cytosol predictions of the 2D DeePiCt CNN^140^ as described above. Results of the CNN-assisted ribosome picking were inspected in tom_chooser^151^ and the remaining, undetected ribosomes were picked manually using e2spt_boxer.py in EMAN2^145^ for up to three rounds.

The coordinates were then used to reconstruct subtomograms and corresponding CTF models in Warp^146^ 1.0.9 with unbinned pixel size, a box size of 140 x 140 x 140 voxel and a diameter of 350 Å. Subtomograms were 3D-aligned and classified in RELION^159^ 3.0.7 into a single class using a low-pass filtered (60 Å) *S. cerevisiae* ribosome map (EMDB 3228) as an initial reference. The resulting average was used as the new reference for the subsequent 3D refinements. Subsequently, we performed refinements in M^147^ with three sub-iterations, allocating 60% of the available resolution to the first sub-iteration. Particles and CTF models were refined by optimizing the particle poses and accounting for non-linear deformations. M-refined subtomograms and CTF models were reconstructed and used for hierarchical 3D classifications into two classes with 25 iterations per job in RELION^159^. The resolution of the resulting averages was calculated based on Fourier shell correlations (FSCs) between the two independently refined half maps. Averages were filtered to the determined resolutions (FSC cut-off criterion=0.143) and visualized either in Chimera^149^ 1.13.1 or ChimeraX^150^ 1.1.1. Focused classifications with soft-edged masks in the PTC and at factor binding sites were conducted in RELION^159^.

In order to determine effective ribosome concentrations, the 3D CNN for ribosome detection was retrained for improved picking efficiency on 29 manually curated tomograms (19 *S. pombe* native state tomograms of which 10 were acquired with defocus only and 9 with the VPP, 5 *S. pombe* 1h ED tomograms acquired with defocus only, 2 *S. cerevisiae* native state tomograms acquired with defocus only and 3 *S. cerevisiae* 1h ED tomograms acquired with defocus only), and validated on one *S. pombe* 1h ED tomograms acquired with the VPP and one *S. cerevisiae* native state tomogram acquired with defocus only, resulting in 0.649 auPRC and 0.4646 auPRC after 150 epochs of training, respectively. The retrained model was first used to predict ribosome localizations on a subset of 7 native state and 4 1h ED tomograms of *S. pombe,* as well as 6 *S. cerevisiae* native state and 6 1h ED tomograms. Based on the predictions, false positives were removed and missing particles added in EMAN2^145^ as described earlier to achieve a 90-95% coverage of ribosome annotations in the data. We next generated segmentations for the cytosol and organelles as described above, and subtracted the organelle volumes from the cytosolic volumes. In the ED tomograms we further segmented the large molecular assemblies. Finally, we determined the ribosome concentrations by dividing the number of ribosomes by the respective cytosolic volume. Since the ribosome concentrations were determined for the same subset of tomograms using CNN predictions only or with additional manual curation, we validated that the results were comparable (Figure S5A). Therefore, we used automated CNN predictions to determine ribosome concentrations on all acquired tomograms for native state cells and at different time points of ED (Table 1, Figure 3B). Of note, the 2D CNN occasionally assigned the nucleus to the cytosolic fraction, which resulted in wrong concentration estimates. Such tomograms (*S.p.* native state: 9/47; *S.p.* 1h ED: 34/117; *S.c.* native state: 15/35; *S.c.* 1h ED: 8/50) were excluded from the analysis based on visual inspection of all cytosol segmentations. For the ectopically acidified *S. pombe* cells, we compared ribosome concentrations determined based on 3 manually curated tomograms and compared them to those determined based on 3D CNN predictions. With an average of 12,320 ± 1373 ribosomes/µm^3^ determined for the manually annotated tomograms and an average of 10,038 ± 1387 ribosomes/µm^3^ determined based on the 3D CNN prediction, there was no significant difference (Kolmogorov-Smirnov test, p=0.6). Thus, automated CNN predictions were used to determine the ribosome concentrations for the tomograms of ectopically acidified *S. pombe* cells (Table 1, Figure 5A). Tomograms of cells under hyperosmolar conditions visually exhibited significant changes in crowding. Despite retraining the 3D CNN using 8 manually curated tomograms of *S. cerevisiae* as input, automated 3D CNN detection failed to reproduce the results in the manual picking (37,811 ± 11,625 ribosomes/µm^3^ from manual curation vs. 11,737 ± 6886 ribosomes/µm^3^ from CNN prediction). Therefore, for tomograms of osmotically stressed cells, only the manually curated data was used to determine the effective ribosome concentrations.

### Polysome fractions

Accurate estimation of polysome fractions requires as complete as possible coverage of ribosome localizations and accurate determination of each ribosome pose. For this reason, we used the particle localizations of the manually curated tomograms described above for the ribosome concentration, in which 90-95% of ribosomes were identified (*S.p.* 7 native state and 4 1h ED*; S.c.* 6 native state and 6 1h ED). We generated subtomogram averages for these ribosomes (following the procedures described above) to refine their positions and orientations. The average of native state *S. pombe* was generated based on 25,901 particles from the 7 manually curated tomograms. The average for the 1h ED *S. pombe* on which the exit and entry site were determined was generated based on 46,120 particles determined by manual curation and CNN prediction. For the subsequent polysome analysis, only the 13,422 particles from the 4 manually curated tomograms were used. For *S. cerevisiae* native state and 1h ED cells, 2x binned subtomograms with a pixel size of 6.85Å and their corresponding CTF models were reconstructed in Warp at a box size of 96 x 96 x 96 voxel and a particle diameter of 300 Å. 27,584 particles for native state and 33,981 particles for 1h ED, were reconstructed. 3D alignment and classification of the subtomograms were carried out in RELION 4.0.11^160^ using a *S. cerevisiae* ribosome map (EMDB 3228, scaled to the data and low-pass filtered to 60 Å) as a reference. The averages were refined over multiple rounds in RELION. The resolution of the averages was determined based on FSC (cut-off criterion 0.143) between the two independently refined half maps. The resolution estimate was 10.7 Å and 10.9 Å for the unbinned (3.3702 Å/px) averages from *S. pombe* native state and 1h ED cells, respectively. For *S. cerevisiae*, the resolution of the 2x binned (6.85 Å/px) native state average was estimated to be 16.2 Å and the average for the 1h ED cells was estimated to 13.5 Å. All averages were visualized in ChimeraX^150^ 1.1.1.

Polysome fractions were determined by assigning ribosomes as monosomes or polysomes using a script in MATLAB 2019a^105^. For this assignment, the positions of mRNA entry and exit sites were determined in Chimera^149^ or ChimeraX^150^ 1.1.1 on ribosome averages of the respective condition. Distances between the mRNA exit site of each ribosome to the mRNA entry site of all neighboring ribosomes were calculated (Figure S6). The distance histograms were fitted with a gaussian distribution. Based on these fits, a distance threshold of 8 nm was chosen as it best described the data. Ribosomes with an mRNA exit site distance of less than 8 nm to the mRNA site of their nearest neighbor were assigned to a polysome. Of note, this approach only identifies polysomes with tightly packed ribosome arrangements, and therefore underestimates polysome numbers that are possible to derive with biochemical methods^46^. Each polysome was assigned a unique identifier and ribosomes within said polysomes were given sequential numbers to determine polysome lengths.

### FAS localization and subtomogram averaging

FAS was localized in 10 tomograms of native state and 16 tomograms of 1h ED *S. pombe* cells. For *S. cerevisiae*, FAS was localized in 3 tomograms of native state cells and one tomogram of 1h ED cells. For all conditions, the complex was manually localized in 4x binned Gaussian-filtered (sigma=3) tomograms using e2spt_boxer.py EMAN2^145^. In *S. pombe* native state cells, a subset of the complexes was additionally localized with a pretrained DeePiCt 3D CNN^140^. The centroids of particles were determined from clusters with a minimal size of 500 voxels and false positives outside the cytosolic volume were removed by multiplication with a cytosol mask considering a contact size of 10 pixel.

Particle coordinates were then used to reconstruct subtomograms and corresponding CTF models in Warp^146^ with unbinned pixel size, a box size of 160 x 160 x 160 voxel and a diameter of 400 Å. FAS particles from each of the conditions in each of the species (native state/1h ED in *S. pombe/S. cerevisiae*) were individually processed by 3D alignment and classification in RELION^159^ 3.0.7. Particles were aligned to the published *S. cerevisiae* map (EMBD 1623^85^). The initial reference was rescaled to the respective unbinned pixel size and low-pass filtered to 60 Å. The resulting averages were refined by applying D3 symmetry. Optimized particles and CTF models were reconstructed following M^147^ refinements and used for hierarchical 3D classifications into two classes in RELION^159^ 3.0.7 with 25 iterations per job, with no major differences between the resulting classes. Resolutions were determined from calculating FSCs between two independently refined half-maps and used to filter the averages (FSC cut-off criterion=0.143). Reconstructed densities were visualized in Chimera^149^ 1.13.1 or ChimeraX^150^ 1.1.1.

The surface charge/electrostatic potential representation of FAS at pH 4.5 and pH 7.5 using the published *S. cerevisiae* atomic model (PDB 2uv8^53^) were generated using a three-step workflow^161^: Pdb2pqr was used to assign protonation states at the specified pH using the default PARSE force field, followed by calculation of the electrostatic potential with the Adaptive Poisson-Boltzmann Solver (APBS) software^162^. The resulting charge maps were visualized through color gradients in ChimeraX^150^.

### Confocal light microscopy and analysis of FAS foci

Exponentially growing *S. pombe* and *S. cerevisiae* cells (OD600=0.3) expressing FAS-mCherry were harvested and either washed thrice with synthetic medium without dextrose containing 20 mM D-glucose or were stress-treated as described above. 10 µM of the pH-sensitive dye CDCFDA was added to visualize cellular environments with a pH below 4.5^163^. 300-400 µl of the cell suspension was transferred to Concanavalin A coated 8-well chambers for imaging. For coating, 1 mg/ml solution of Concanavalin A in water was applied to the bottom of the imaging chamber to cover it evenly, and incubated for 30 minutes. Then, the Concanavalin A solution was aspirated and the imaging chamber was washed thrice with distilled water prior to addition of the yeast cell suspension. Before imaging, cells were allowed to attach to the Concanavalin A-coated surface. A total of 18 z-slices with a spacing of 640 nm and an image size of 792 x 792 pixels at a pixel size of 85.2 nm were acquired with the Zeiss Zen Software on an LSM780 or LSM880 inverted microscope using the 63x/1.40 NA oil immersion objective with Immersol 518F at 30°C with a 561 nm DPSS and a 488 nm argon excitation laser.

The number of FAS foci per cell was determined in two ways: 1) using an automated analysis pipeline that employs Fiji^153^, ilastik^152^, R (version 3.6.3 (R Core Team 2013)) and RStudio version 1.2.5033^164^, or 2) by 2D cell detection with YeaZ^165^, manual thresholding for binarization and separation of fused cells using the watershed algorithm, followed by a two-step particle analysis in Fiji^153^.

The first approach was used to determine the number of FAS foci in the timelapse series comparing ED and control cells (Figure 2). Here, the central z-slice was determined in two consecutive steps in Fiji^153^: 1) The position of an initial central z-slice was selected based on the highest standard deviation of fluorescence intensity signal. 2) This position was then refined by identifying the z-slice with the lowest integrated intensity in the bright field channel within the range of two slices above and below the initially selected z-slice using an IsoData threshold algorithm^166^. The number of foci was then determined using the 3D object counter in Fiji^167^ within four slices above and below the central z-slice. The minimum voxel size for object detection was set to 9 at an empirically determined threshold of 132 plus the average FAS-mCherry intensity. Individual cells were segmented in ilastik^152^. Maximum intensity projections of the three most central z-slices were subjected to pixel classification. Cell wall and background were detected by a network trained on manual segmentations with selected features of Gaussian smoothing, Laplacian of Gaussian, Gaussian gradient magnitude, difference of Gaussians, structure tensor eigenvalues and Hessian of Gaussian eigenvalues to generate probability maps as input for boundary-based cell segmentations. A watershed algorithm with empirically determined thresholds of 0.2 to 0.3 and pre-smoothing of 1.0-2.0 was applied. Minimum boundary and superpixel sizes were set to 0. To iteratively select or exclude classified segmentation edges, a network was trained with clustered seed labeling and thin structure preservation. Next, the Nifty FMGreedy solver with a bias parameter of 0.4 was used to generate an initial multicut segmentation. The latter, together with the brightfield z-projections, were then used as input for object classification to generate binary masks of individual cell segmentations. The binary masks were multiplied with the multicut segmentations in R to assign zero intensity values for the background and individual integers for each cell. This result was then integrated with the foci counts from the 3D object counter to track the number of foci per cell in control and ED cells over time.

The second approach was used to determine the number of FAS foci under different stress conditions, while monitoring the pH. Individual cells were segmented on manually selected center slices using YeaZ^165^. In Fiji^153^, the output masks where thresholded to generate binary masks and subsequently assign ROIs to the individual cells through the analyze particles tool. To determine the number of FAS foci per cell, the mCherry channel was thresholded based on the intensity of dispersed FAS in the corresponding control cells and Fiji’s analyze particles tool was applied on a per cell basis to detect the individual foci. Cell volumes were determined based on brightfield center slices of confocal microscopy stacks. Therefore, cells were segmented using YeaZ^165^ resulting in binary outputs that were used to automatically reconstruct cell volumes with the Pomegranate tool^168^. This approach was chosen since stressed cells moved substantially between frames despite the attempt to immobilize them with Concanavalin A, which prevented reasonable volume quantification based on stacks. Raw images of the fluorescent channels were used for quantification and adjusted for brightness/contrast and smoothed in ImageJ/Fiji^153^ for visualization. For quantitative analysis, the number of processed cells (n) from single experiments is indicated with the data.

### FRAP measurements of FAS-mCherry

FRAP experiments were performed on an LSM 780, using either a C-Apochromat 40x/1.2 NA W Korr FCS water objective or a Plan-Apochromat 63x/1.4 NA Oil DIC objective. *S. pombe* FAS2-mCherry or *S. cerevisiae* FAS1-mCherry cells were imaged at an OD600 of 0.1 with the following imaging and FRAP conditions:

*S. pombe* FAS2-mCherry: Imaging: 500 images were collected at an interval of 1 second from an image size of 140 x 120 pixels (117 nm pixel size) using 2% laser power (λ_exc_ = 561 nm, DPSS laser) with 2.46 µs pixel dwell time and a 0.1 second frame time; Fluorescence bleaching and recovery: a rectangular area of 44 x 16 pixels was bleached once after image number 40 with 100% laser power and a pixel dwell time of 7.49 µs. The fluorescence recovery after photobleaching was measured for 5 cells per condition.

*S. cerevisiae* FAS1-mCherry: Imaging: 300 images were collected at an interval of 0.1 seconds from an image size of 132 x 132 pixels (85.2 nm pixel size) using 15% laser power (λ_exc_ = 561 nm, DPSS laser) with 1.65 µs pixel dwell time and a pixel dwell time of 0.07 seconds; Fluorescence bleaching and recovery: a rectangular area of 63 x 16 pixels was bleached once after image 20 with 40% laser power and a pixel dwell time of 3.3 µs. Fluorescence recovery after photobleaching was measured for 21 cells for each of the normal nutrient control and the energy depletion conditions.

FRAPAnalyser^154^ version 2.1.0 (https://github.com/ssgpers/FRAPAnalyser) was used to fit recovery curves. The data was double-normalized against the background and either a whole cell reference for the larger *S. pombe* or a rectangular area the size of the bleaching area for *S. cerevisiae*, which covered parts of a neighbouring, unbleached cell using the following equation:

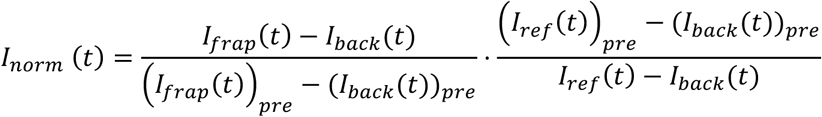

The normalized data was fitted using the Levenberg-Marquardt method with a single (control cells) or a double (energy depletion) exponential model with the following equations:

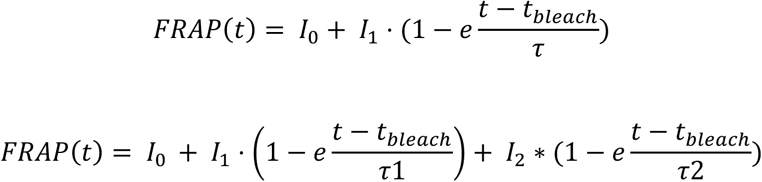

Recovery half-times were calculated as *t*_1/2_ = *τ* ⋅ *ln*2 and immobile fractions determined from (1 − *I*_0_ − *I*_1_)/(1 − *I*_0_). We calculated two separate recovery half times (t_1/2, 1_ and t_1/2, 2_) and immobile fractions (I_immo, ½, 1 and 2_) for the two populations of tagged FAS (free FAS and FAS assembled into foci) from a single bleaching area since the foci are smaller than the minimal bleaching area.

## QUANTIFICATION AND STATISTICAL ANALYSIS

The details of the quantification and all statistical analyses are included in the relevant figure captions or sections of the Method Details. all statistical tests were conducted using the scipy python package^169^.

---

**Figure S1.**
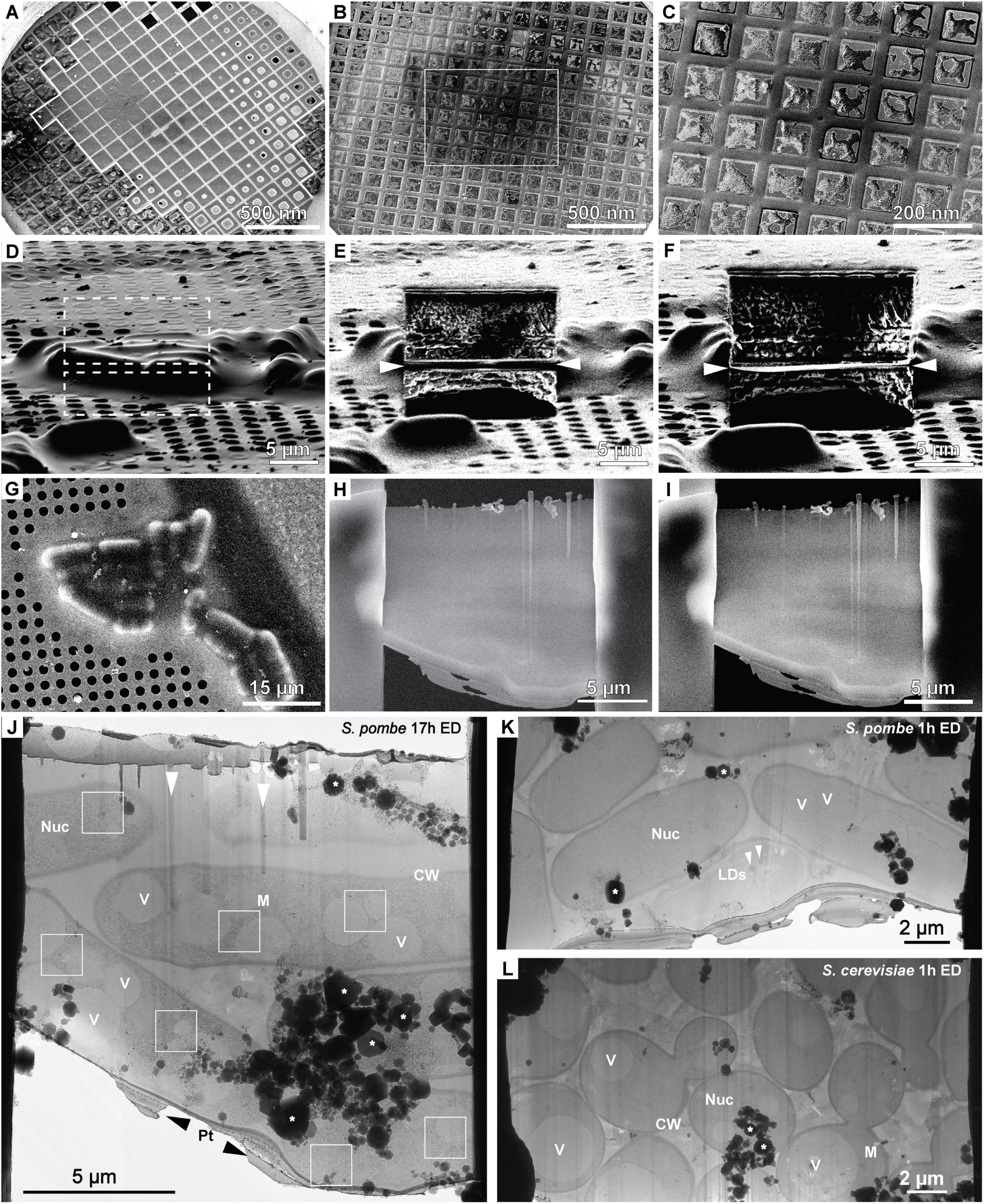
Cryo-FIB preparations of yeast cells for cryo-ET, related to Figure 1 and Table 1. (A-C) Cryo-SEM images of yeast cells plunge-frozen on transmission electron microscopy (TEM) grids: (A) A gradient of *S. pombe* cells across a grid plunge-frozen at OD600 0.6 with a Vitrobot (Thermo Fisher Scientific). Area with empty squares at the center of the grid is outlined. (B) *S. pombe* cells frozen at OD600 0.3 with a Leica GP plunger. (C) Magnified view of boxed area in (B) shows homogenous distribution of cells and optimal ice thickness inside grid squares at the grid center. (D, G) *S. pombe* cells targeted for cryo-FIB milling viewed from shallow angle in the FIB view (D) and top view in the SEM (G). Milling patterns are represented as white dashed boxes in D. (E-I) FIB (E, F) and SEM views (H, I) of cells targeted in (D, G) milled to 1 µm thickness (E, H), and of the lamella at final thickness of approximately 200 nm (F, I). (J) Cryo-TEM overview of lamella in (I) showing six *S. pombe* cells at 17h ED. Cell wall (CW), vacuoles (V), mitochondria (M) and nuclei (Nuc) are annotated. Seven positions of cryo-ET acquisition are highlighted by white boxes. Curtaining artifacts from slower ablation of lipid droplets (white arrowheads), ice contaminants introduced during transfer into the TEM (*) and the protective organometallic platinum layer at the lamella front (Pt, black arrowheads) are indicated. (K, L) Cryo-TEM lamella maps of *S. pombe* (K) and *S. cerevisiae* (L) at 1h ED. LDs: lipid droplets (white arrowheads), Nuc: nucleus, CW: cell wall, M: mitochondrion. Asterisks indicate ice contamination.

**Figure S2.**
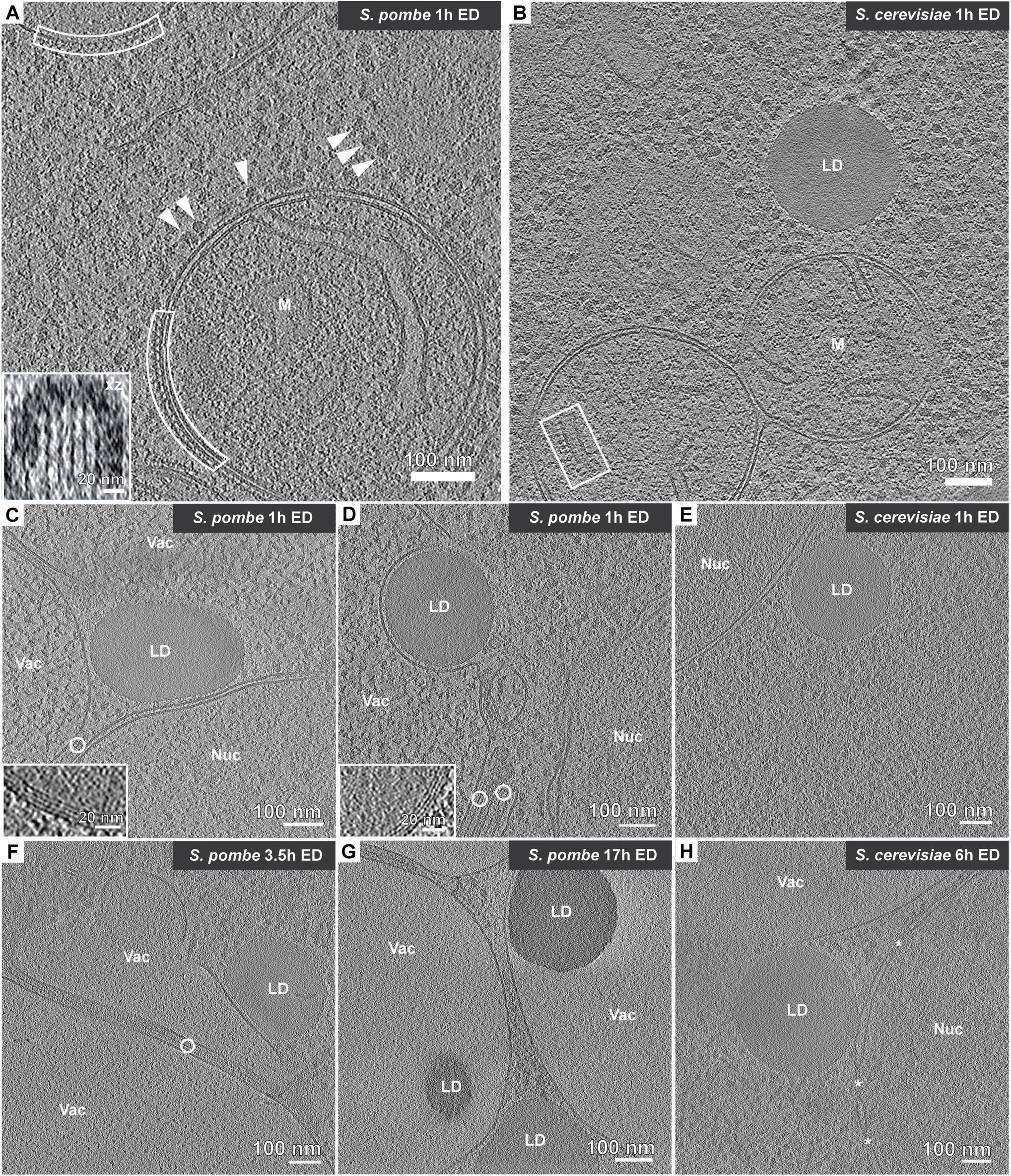
Energy depletion induced changes to organelle morphologies, related to Figure 1. (A, B) Tomographic slices of cells at 1h ED showing spherical mitochondria morphologies. (A) In *S. pombe*, ribosomes (arrowheads) and surface lattices were observed at the outer membrane (white boxes) of a subset of mitochondria. Inset: Enlarged xz-view of a mitochondrial-surface lattice. The lattices were not observed in *S. cerevisiae*. (B) ATP synthases (white box) decorate the cristae of an *S. cerevisiae* mitochondrion undergoing fission. (C-H) Changes in lipid droplet (LD) morphology under ED. (C, D) In *S. pombe* at 1h ED, LDs with crystalline outer layers^76,77^ (enlarged in insets) appear deformed in vicinity of the vacuole and nucleus (C), similar to recent observations in glucose restricted cells^77^, or are partially engulfed by the vacuole (D). Monolayers of FAS (white circles) staked on the surface of the vacuoles. (E, H) In *S. cerevisiae*, at 1h (E) and 6h (H) ED, LDs remain spherical and located in proximity to the nucleus. (F) In *S. pombe* at 3.5h ED, a LD is deformed by contact with a vacuole that is fractionated by a FAS monolayer array (white circle). (G) At 17h ED, LDs are taken up by vacuoles. M: mitochondrion, LD: lipid droplet, Vac: vacuole, Nuc: nucleus, white circles: FAS, asterisks: nuclear pores.

**Figure S3.**
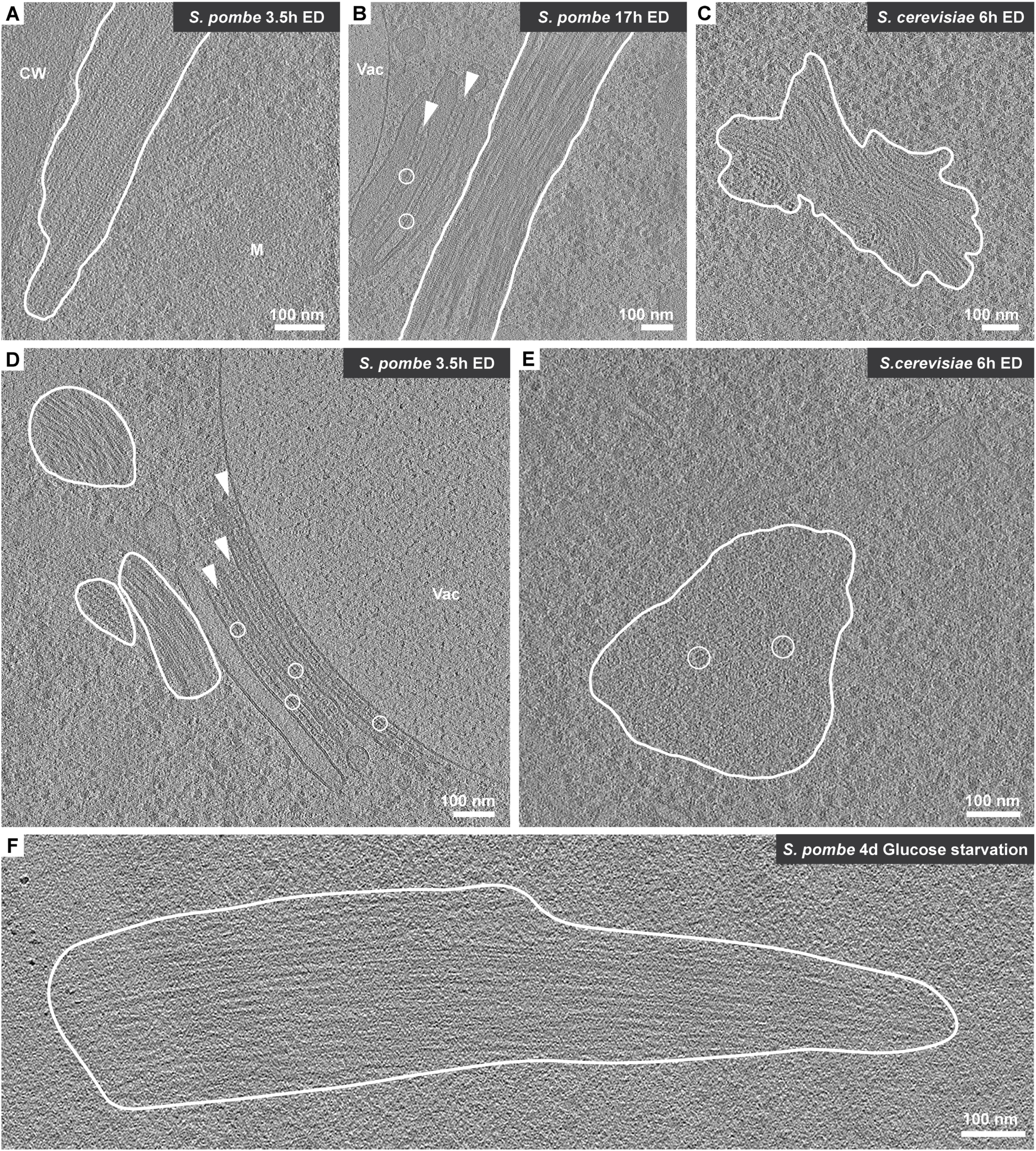
Species-dependent assemblies grow beyond one hour of energy depletion, related to Fig. 1. Macromolecular assemblies (encircles by white lines) occupy large volume fractions at late timepoints of ED. (A, B) Lattices observed at 1h ED (Figure 1H) occupied larger cytosolic volume fraction at 3.5h (A) and 17h (B) ED in comparison to 1h ED in *S. pombe*. (B) Assemblies at 17h observed in proximity to FAS (white circles) arrays stacked between membrane (white arrowheads). (C) Polymeric assemblies observed in 1h ED *S. cerevisiae* (Figure 1I) occupied a larger volume fraction at 6h of ED. (D) FAS (white circles) arrays arranged between several membrane stacks (white arrowheads) at 3.5h ED in *S. pombe* observed to co-exist with three different types of filament assemblies. (E) At 6h ED compared to 1h ED in *S. cerevisiae*, FAS (encircled in white) concentrated in larger areas of the cytosol. (F) Large macromolecular assemblies of similar morphology as those observed under ED were detected after four days of glucose starvation. M: mitochondrion, CW: cell wall, Vac: vacuole.

**Figure S4.**
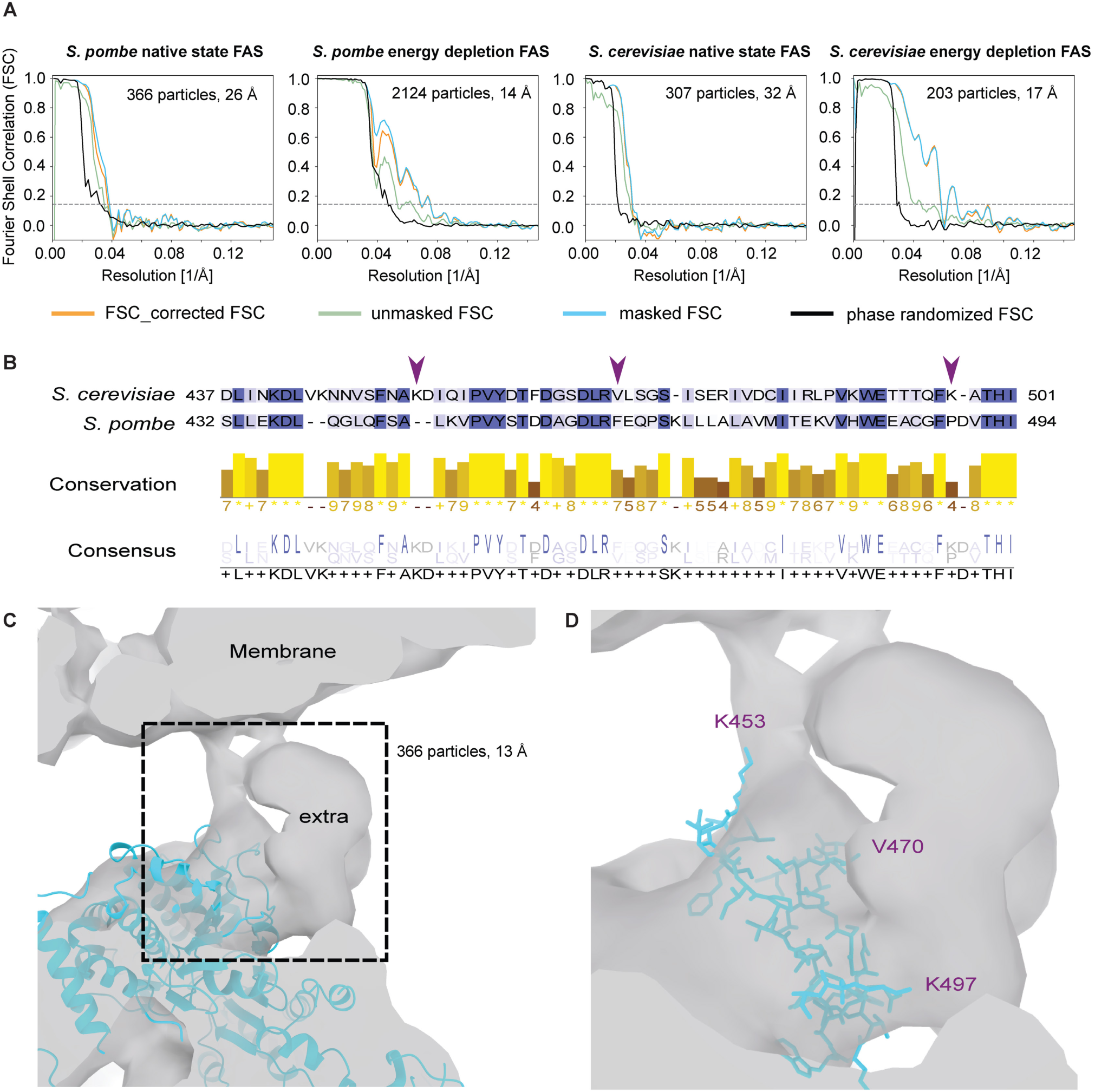
FAS subtomogram averaging and sequence alignment, related to Figure 2. (A) Fourier shell correlation (FSC) curves of FAS subtomogram averages determined with D3 symmetry of the well-aligned FAS classes in native state (all particles from 10 tomograms for *S. pombe* and 53% of localized particles from 16 tomograms for *S. cerevisiae*) and 1h ED (99% of particles from three tomograms for *S. pombe* and 59% of particles from one tomogram for *S. cerevisiae*) with particle numbers corresponding to those in Figure 2A. FSC threshold of 0.143 shown as dotted lines^166^, with the determined resolution indicated. (B) Sequence alignment of FAS1/ß for *S. cerevisiae* and *S. pombe* at the interaction interface depicted in C. Top: residues colored by sequence conservation (high: dark blue, low: white); middle: conservation based on physicochemical properties determined using the AMAS method in Jalview^167^; bottom: consensus highlighting identical residues (large letters). Purple arrowheads highlight residues labeled in D. (C) Zoomed in view of the subtomogram average of *S. pombe* FAS at 1h ED showing its interface with the membrane (grey) fitted with the crystal structure of the FAS1/ß unit of *S. cerevisiae* (PDB: 2uv8^80^, cyan). Two loops of the FAS structure face the extra density connecting to the adjacent membrane. (D) Enlarged view into the area highlighted in C (dashed box) with *S. cerevisiae* FAS interface loops represented as sticks.

**Figure S5.**
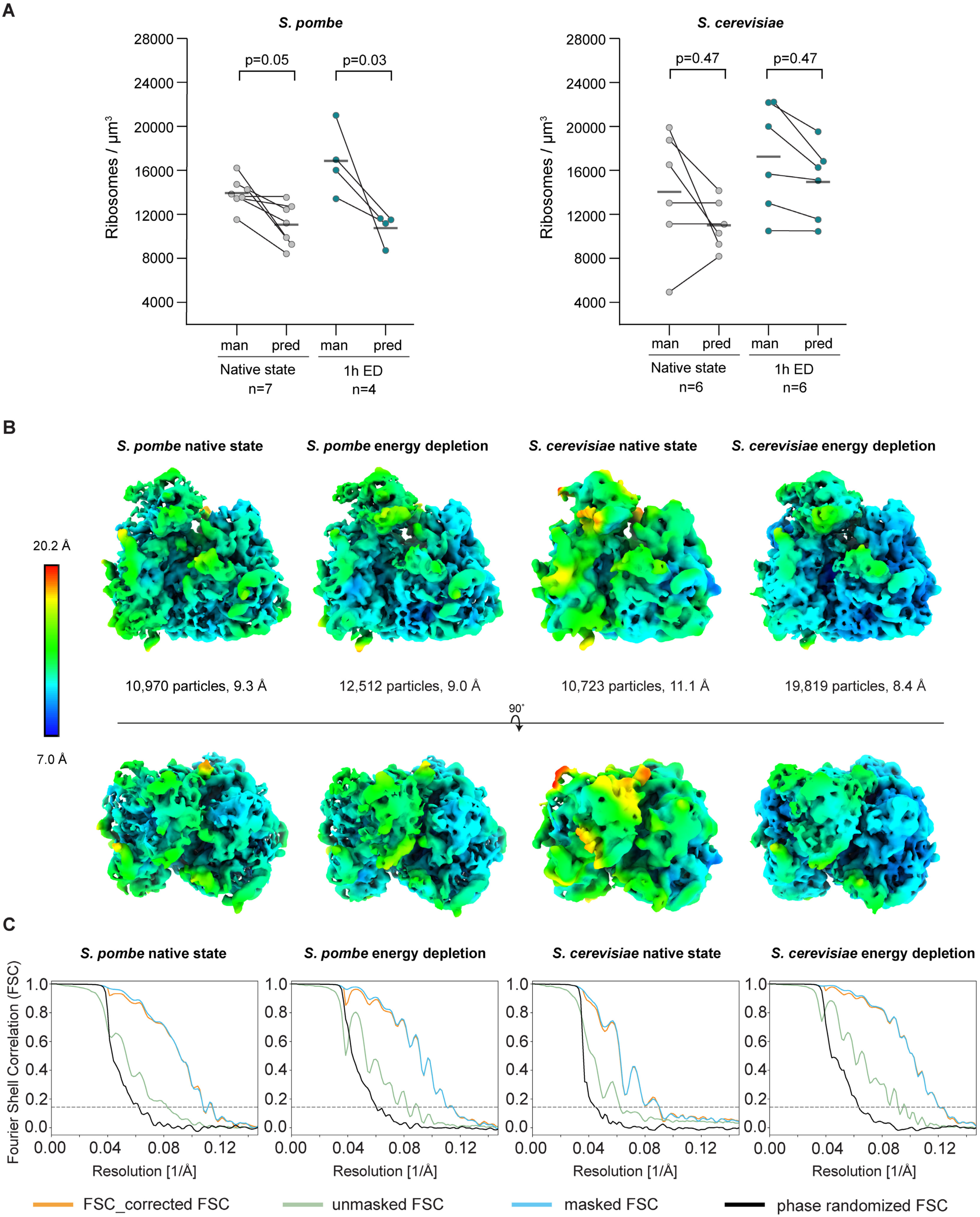
Analysis of ribosome concentrations and subtomograms, related to Figure 3. (A) Comparison of ribosome concentrations determined in the same tomograms of native state and 1h ED cells based on manual curation (man) versus automated detection with supervised 3D neural networks (pred)) in *S. pombe* and *S. cerevisiae* connected by lines. Number of tomograms analysed (n) is indicated for each condition. The benchmarking shows that automated detection by the trained 3D CNN provides similar performance to manual expert curation. (B) Local resolution maps of well-aligned ribosome classes from native state (42% and 47% of particles from 10 and nine tomograms of all localized particles in *S. pombe* and *S. cerevisiae,* respectively) and 1h ED cells (27% and 36% of all localized particles in 20 *S. pombe* and 16 *S. cerevisiae* tomograms). Particle numbers and resolution are provided for each of the maps. Top row: side view, bottom row: top view. (C) FSC curves of the well-aligned ribosome subclasses refined in M^142^. FSC threshold of 0.143 indicated as dotted lines^166^ was used to determine the map resolution indicated in B.

**Figure S6.**
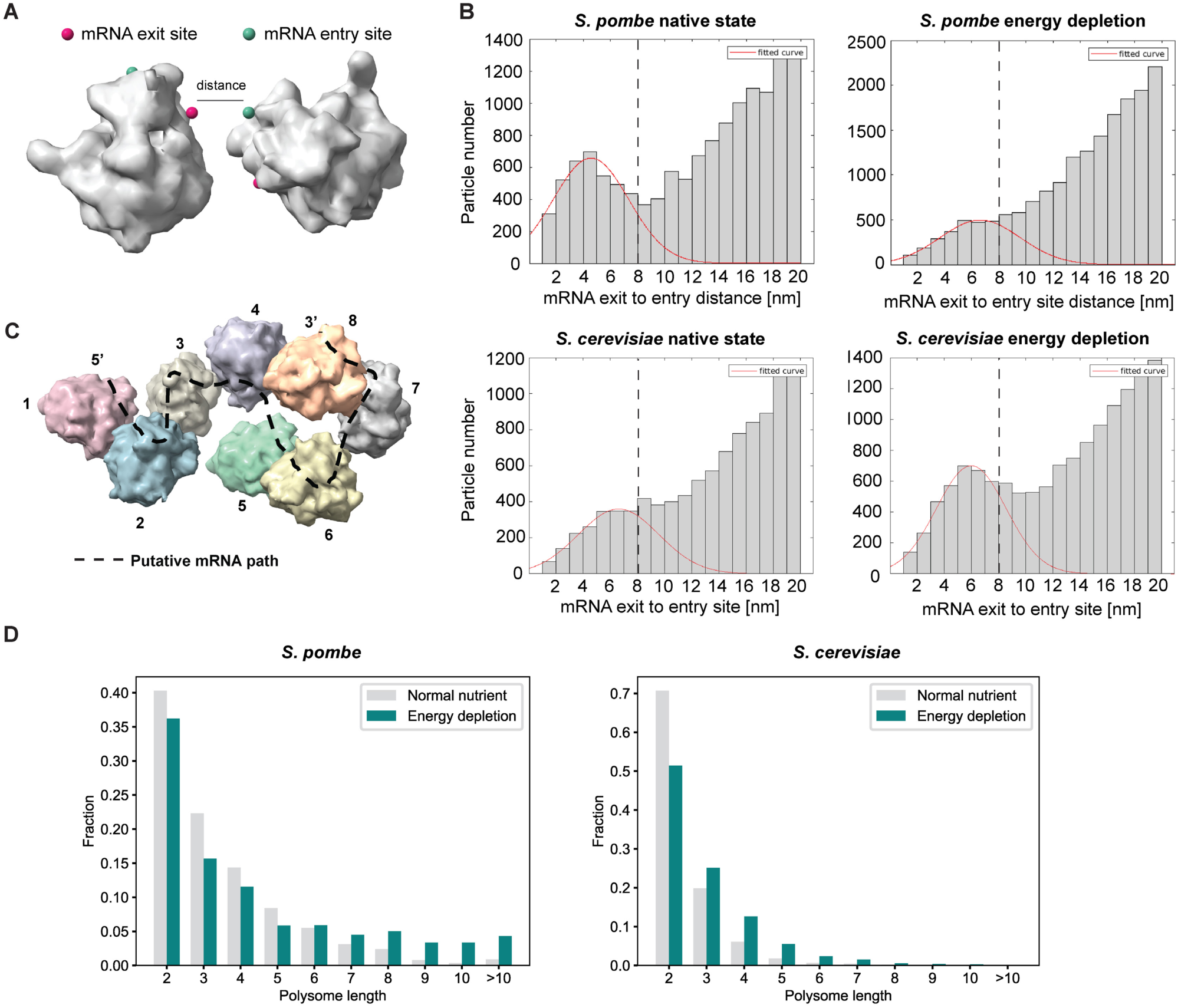
Polysome analysis, related to Figure 3. (A) The distance between the mRNA exit site of one ribosome and the mRNA entry site of the closest neighbouring ribosome was used to detect polysomes in the data. (B) mRNA exit-to-entry site distance distribution for native state and 1h ED *S. pombe* and *S. cerevisiae* cells. Gaussian fits (red fitted curve) to the polysome peak indicate a distance threshold of 8 nm, which was used to assign individual ribosomes to the same polysome. (C) Representative long polysome consisting of 8 ribosomes with the corresponding putative mRNA path indicated (dashed line). (D) Distribution of polysome lengths in normal nutrient state and 1h ED *S. pombe* and *S. cerevisiae*.

**Figure S7.**
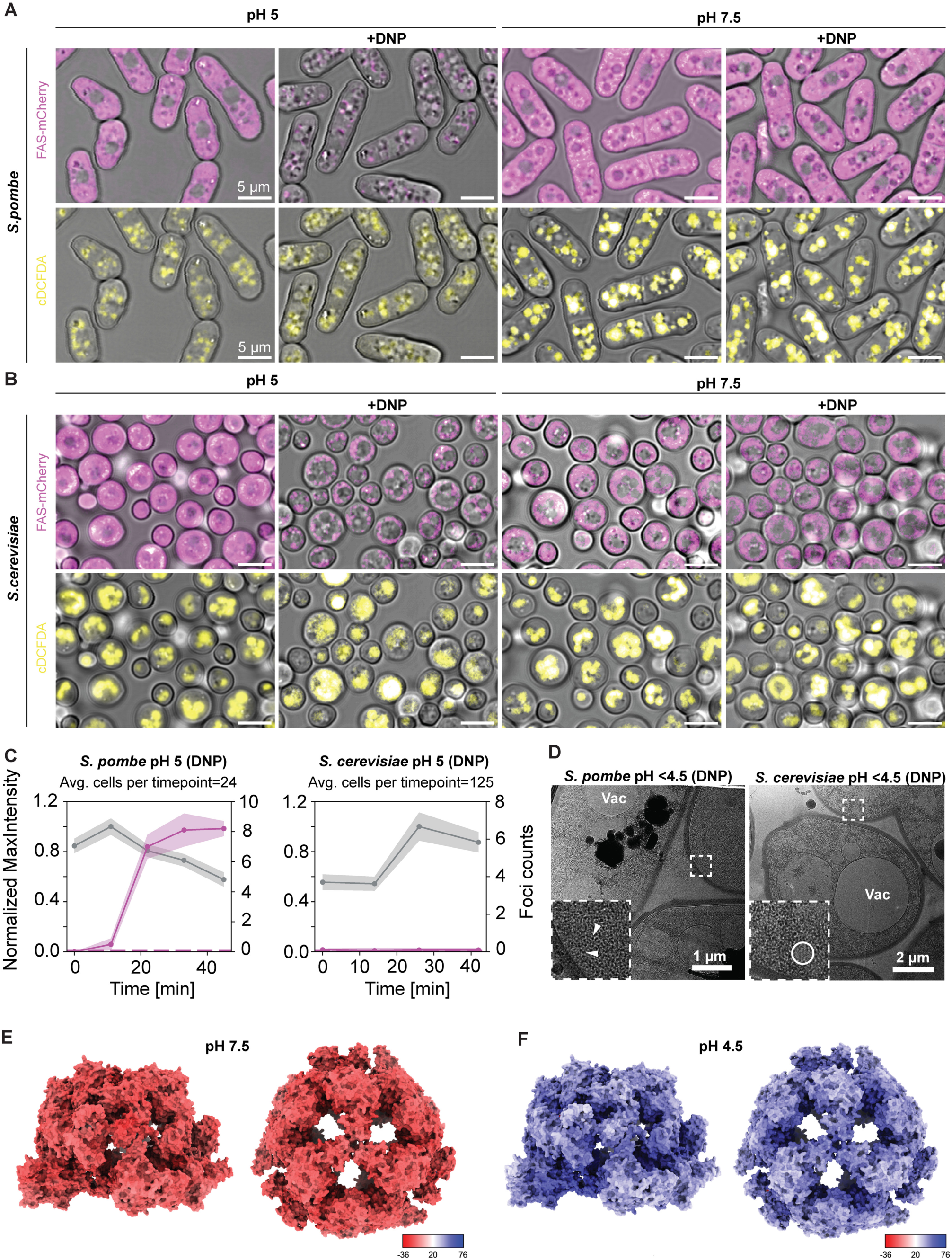
Ectopically induced cytosolic acidification triggers FAS assembly, related to Figure 4 and 5. (A, B) Brightfield and confocal microscopy stacks of live *S. pombe* (A) and In *S. cerevisiae* (B) cells expressing FAS-mCherry, co-stained with the pH indicator cDCFDA in different pH media. FAS formed foci in cells equilibrated to pH 5 through addition of proton carrier (2,4-Dinitrophenol; +DNP) and pH titration of the cell growth medium. In the absence of the proton carrier and when equilibrated to pH 7.5 (right, +DNP), FAS remained dispersed. (C) Quantification of FAS foci (magenta, right y-axis) in the cytosol of *S. pombe* and *S. cerevisiae* (pH 5, +DNP), plotted alongside the maximum intensity of the cDCFDA signal (grey, left y-axis) over time. Not all *S. cerevisiae* cells reacted to this condition, similar to the response observed in unbuffered medium with DNP, leading to low foci counts (Figure 4C). Dotted lines: starting levels of FAS foci counts. (D) cryo-TEM images of *S. pombe* after 50 min and of *S. cerevisiae* after 38 min in acidic medium (pH <4.5, + 2,4-DNP). The cytosol of *S. pombe* shows discernible ribosomes (inset: arrowhead), while the cytosol of *S. cerevisiae* appears aggregated leading to large patches that seem devoid of ribosomes (inset: circle). Vac: vacuole. (E, F) Surface charge/electrostatic potential representation of FAS (PDB 2uv8^80^, Methods) at pH 7.5 (E) and pH 4.5 (F).

## Notes

### Competing Interest Statement

The authors have declared no competing interest.

